# Parietal alpha stimulation causally enhances attentional information coding in evoked and oscillatory activity

**DOI:** 10.1101/2023.11.14.567111

**Authors:** Runhao Lu, Elizabeth Michael, Catriona L. Scrivener, Jade B. Jackson, John Duncan, Alexandra Woolgar

## Abstract

Selective attention is a fundamental cognitive mechanism that allows people to prioritise task-relevant information while ignoring irrelevant information. Previous research has suggested key roles of parietal evoked potentials and alpha oscillatory responses in spatial attention tasks. However, the informational content of these signals is less clear, and their causal effects on the coding of multiple task elements are yet unresolved. Here, we used concurrent TMS-EEG to causally manipulate parietal alpha power and evoked potentials and investigate their roles in coding multiple task features (*where to attend*, *what to attend to*, and *visual stimulus*) in a selective attention task. First, using EEG-only data, we found that evoked potentials coded all three types of task-relevant information with distinct temporal dynamics, and alpha oscillations carried information regarding both where to attend and what to attend to. Then, we applied rhythmic-TMS (rTMS) at individual alpha frequency over the right intraparietal sulcus (IPS), while concurrently measuring EEG. Compared with control arrhythmic-TMS, alpha rTMS increased alpha power and inter-trial phase coherence and yielded more negative posterior-contralateral evoked potentials. Moreover, alpha rTMS causally and specifically improved multivariate decoding of the information about *where to attend* (but not *what to attend to* or *feature information*) during task performance, with decoding improvements predicting changes in behavioural performance. These findings illuminate the dynamics with which the complementary aspects of a selective attention task are encoded in evoked and oscillatory brain activity. Moreover, they reveal a specific and causal role of IPS-controlled evoked and oscillatory activity in carrying behaviour-driving information about where to focus attention.

## Introduction

Selective attention is a fundamental cognitive mechanism that allows individuals to prioritise relevant information while ignoring distracting and irrelevant information ^1–3^. This selective mechanism can be deployed in various ways depending on factors such as spatial location, task rules, or visual features of an object ^4–7^. It is well documented that both evoked potentials (also known as event-related potentials; ERP) and oscillatory activities are modulated by selective attention ^8–16^. For example, posterior-contralateral ERP components, such as N2pc and sustained posterior contralateral negativity (SPCN), and alpha power (8-13 Hz) are known to track attention allocation to a cued location ^17–23^.

EEG-based multivariate pattern analysis (MVPA) can provide complementary insights into how EEG signals represent various task aspects ^24,25^. There is growing evidence that various task aspects, including cued locations (*where to attend*) ^26,27^, task rules (*what to attend to*) ^26,28^ and *visual feature information* (e.g., stimulus colour, shape, etc) ^26,29,30^ can be decoded from broadband EEG or magnetoencephalography (MEG) signals. However, it remains less clear how different aspects of these EEG signals (e.g., evoked potentials and alpha oscillations) contribute to the representation of these different aspects of information. Intriguingly, one study on working memory, using MVPA with EEG, found that evoked potentials code information about both target location and target features while alpha oscillations only code location information ^8^. Similarly, another recent study found that alpha power, rather than evoked potentials, could decode memory information that was thought to be “activity-silent” ^31^. These findings indicated potential functional dissociations between evoked potentials and alpha oscillations. To date, however, it remains unclear how evoked potentials and alpha code different types of task-relevant information in selective attention tasks.

For visual feature information, a further question concerns how attention modulates the feature representations carried by evoked potentials and/or alpha. There are three possibilities proposed by previous studies on this question: the object-based attention hypothesis ^32–35^, the feature-based attention hypothesis ^6,7,36^, and the multiplicative hypothesis ^26,37^. The object-based attention hypothesis proposes that both task-relevant and task-irrelevant features of an object will be processed if any one feature of an object is selected. In contrast, the feature-based hypothesis suggested that task-relevant features of all the presented objects are attended, while the multiplicative hypothesis suggests that only the task-relevant feature of the attended object is prioritised for in-depth processing.

Beyond observing the correlational relationship between neural signals (e.g., evoked potentials and alpha oscillations) and attentional information coding, combining EEG recordings with non-invasive brain stimulation techniques, such as transcranial magnetic stimulation (TMS), provides a promising way to manipulate oscillatory activities and to examine how they causally influence information coding ^38–41^. TMS entrainment studies have found that repetitive rhythmic TMS pulses in a certain frequency band (e.g., alpha frequency) can entrain (i.e., enhance) local power and phase coherence in that specific frequency ^42–45^. Also, there is evidence that TMS at the parietal cortex can influence amplitudes of attention-related evoked potentials like posterior contralateral components ^46–48^. They together indicate that using TMS at alpha frequency may manipulate alpha oscillations and evoked potentials related to attention, which may further influence task-relevant information coding.

In this two-part study, we first investigated the decodability and temporal dynamics of evoked potentials and alpha power in coding different aspects of selective attention using EEG. Using a selective attention paradigm (Figure 1A) and time-resolved MVPA ^25,49^, we examined the representational dynamics of three task aspects (*where to attend*, *what to attend to*, and *visual feature information*) based on evoked potentials and alpha power separately. Subsequently, we asked how attentional states modulated visual feature coding and how the results aligned with previous hypotheses (i.e., object-based attention, feature-based attention, and multiplicative hypothesis). In the second part of this study, we intervened on pre-stimulus parietal alpha, to observe its causal downstream effect on evoked potentials, post-stimulus alpha power, and coding of different aspects of selective attention. For this, we used concurrent TMS-EEG with either rhythmic-TMS (rTMS) at each individual’s task-based alpha frequency (IAF), or control arrhythmic-TMS (aTMS). We applied this stimulation over the right IPS, one of the regions associated with attentional control, which is thought to regulate attention-related evoked and oscillatory activities ^50–53^. We first checked whether our EEG-only findings replicated in the TMS-EEG data. Next, we examined whether the rTMS protocol entrained alpha power and inter-trial phase coherence (ITPC) relative to aTMS, and whether rTMS affected amplitudes of posterior-contralateral ERPs. Then, we examined the causal effect of alpha rTMS on the three aspects of task information coded by evoked potentials and alpha power. Finally, we asked whether the rTMS-induced changes in information coding were important for behavioural performance, by correlating them to rTMS-induced change in behaviour across subjects.

**Figure 1.**
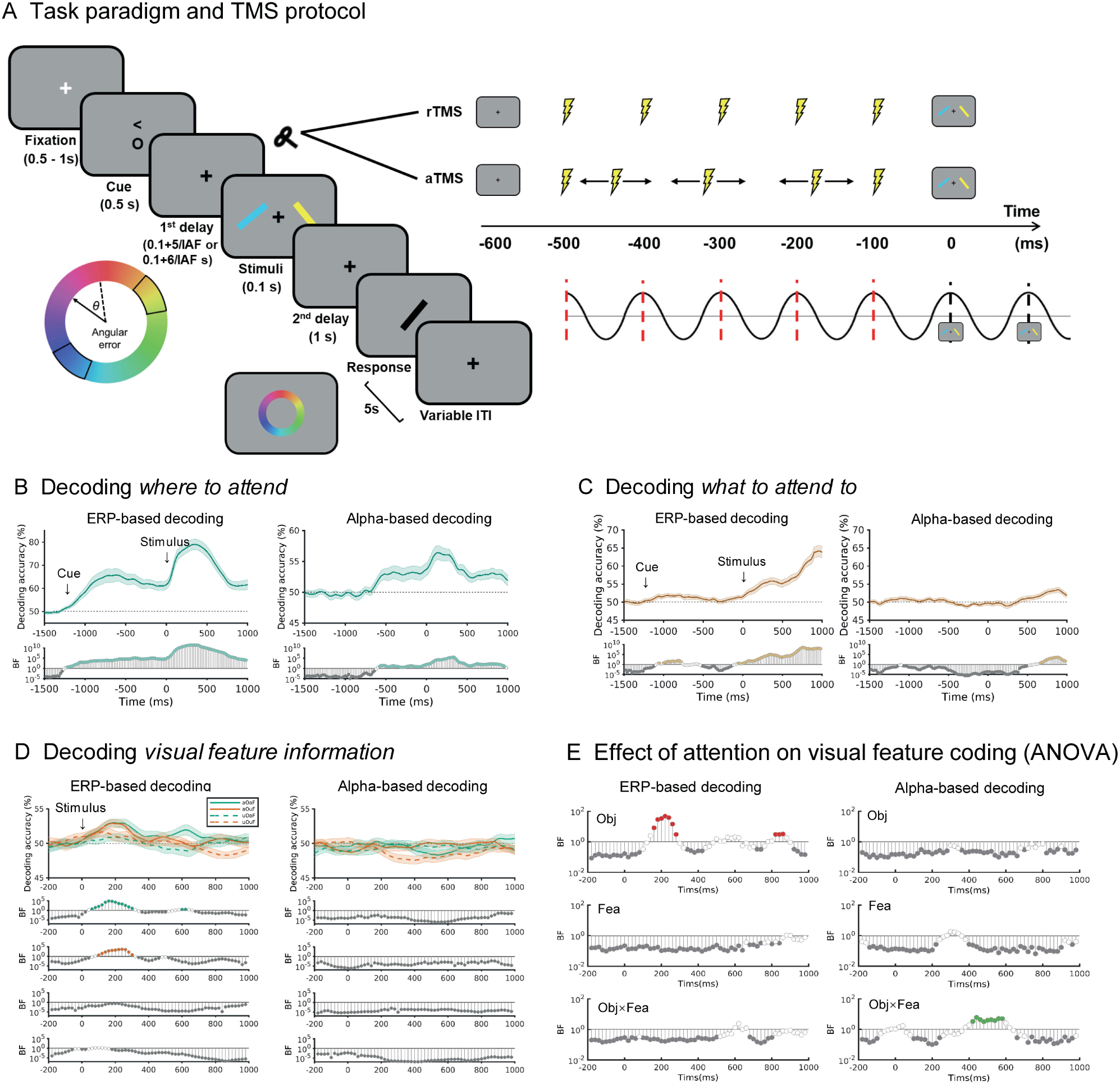
Task paradigm and decoding of *where to attend*, *what to attend to* and *visual features* using EEG-only data. (A) Task paradigm. After a variable fixation, participants were cued to attend to either left or right (‘<’ means attend left; ‘>’ means attend right), and to attend to either the colour or orientation of the attended object (‘c’ means attend colour; ‘o’ means attend orientation). Following the first delay (600 ms or 700 ms on EEG-only trials, adjusted for IAF on TMS-EEG trials), two stimuli (coloured tilted bars) were presented on each side of the screen for 100 ms. After the second delay (1000 ms), participants were asked to make a response using a computer mouse. In the orientation task, participants used the mouse to manipulate the angle of a black bar at the centre of the screen to match the angle of the cued object and then clicked the right mouse button when it was completed. In the colour task, participants clicked the colour of the cued object in a colour wheel. After making the response, there was a variable inter-trial interval (ITI) before the next trial. The response time and the ITI was 5 seconds in total. During TMS-EEG sessions, five-pulse TMS was applied to participants’ right intraparietal sulcus (IPS), beginning 100 ms into the first delay, before stimulus onset. These pulses were either rhythmic (rTMS), or arrhythmic (aTMS). Stimulus onset aligned with the first or the second peak of the TMS-induced cycle (50% of trials each). The interval between pulses, as well as between the last pulse and the stimulus onset, varied depending on participants’ individual alpha frequency (IAF, determined using 3 blocks of EEG-only trials). For EEG-only sessions, no TMS pulses were given. This picture illustrates timings when IAF is 10 Hz and the stimulus is delivered at the peak of the first post-TMS cycle. The time point 0 indicates stimulus onset. (B-D) ERP-based decoding and alpha power-based decoding results of (B) *where to attend* (attend left vs. attend right), (C) *what to attend to* (attend colour vs. attend orientation), and (D) *visual feature information* (yellowish vs. blueish colour, leftward vs. rightward orientation). Time in all plots is relative to stimulus onset (ms). Note the different x-axis scale in (D), in which we only examined coding of visual features from shortly before they were presented. Shaded error bars indicate the standard error over participants. The bottom rows of each plot show the Bayes Factors (BF_10_) at each time point on a log scale, with BF_10_ < 1/3 marked in grey and BF_10_ > 3 highlighted with coloured circles. aOaF: attended feature of attended object; aOuF: attended feature of unattended object; uOaF: unattended feature of attended object; uOuF: unattended feature of attended object. (E) Bayesian ANOVA (BF_10_) examining the effect of feature and object attention on the representation of visual feature information from on ERP (left) and alpha power (right) for each timepoint. Three rows from top to bottom represent the BF for the main effect of object, the main effect of feature, and the interaction between object and feature. Timepoints with BF_10_ < 1/3 are represented by grey circles and BF_10_ > 3 by coloured circles.

## Results

### Part 1: EEG-only data

In the first part of the study, we used EEG-only data to test whether and how evoked potentials and alpha power, separately, code three types of information during a selective attention task: *where to attend* (attend left vs. right), *what to attend to* (attend colour vs. orientation), and *visual feature information* (yellowish vs. blueish colour, leftward vs. rightward orientation). For *visual feature information*, we separated the data into four attention conditions according to the combination of whether the decoded feature (e.g., colour) was relevant for the task (e.g., colour in the colour task, attended feature) or irrelevant for the task (e.g., colour in the shape task), and whether it pertained to the attended object (e.g., object on the left in attend left trials, attended object) or the distractor (e.g., object on the left in attend right trials, unattended object). We compared coding between these conditions to test how selective attention modulated information coding. To isolate the separate representational content of evoked responses and alpha power, we used time-resolved MVPA on whole-brain ERP data that were filtered to reflect evoked potentials (1-6 Hz) or transformed into a moment by moment estimate of alpha power (8-13 Hz), following Bae and Luck ^8^. At each timepoint, we assessed the evidence for information coding using Bayesian analyses ^54^.

### Behavioural results

Participants performed well on the task with a mean error of 9.79 degrees for colour and 6.75 degrees for orientation, and mean response time (RT) of 1.65 seconds and 2.10 seconds respectively (Table S1). 2*2 Bayesian repeated measure ANOVAs (cued location: attend left vs. attend right; cued feature: attend colour vs. attend orientation) on RT and response error showed very strong evidence for the main effect of cued feature on RT (BF_10_ = 7.32×10^19^) and response error (BF_10_ = 9.69×10^9^). This showed that participants had higher response error but shorter RT in the colour task than those in the orientation task. However, we do not interpret these differences, due to the different scales (degrees of colour vs degrees of orientation) and methods of responding in the two tasks. There was insufficient evidence (intermediate Bayes Factors) to determine whether there was a main effect of cued location (BF_10_ = 0.63) or interaction between cued location and cued feature (BF_10_ = 0.65) on response error. There was evidence for the null hypothesis that there was no main effect of cued location (BF_10_ = 0.12) and no interaction between cued location and cued feature (BF_10_ = 0.16) on RT.

### Representational content of evoked potentials and alpha oscillatory responses

As shown in Figure 1B, information about *where to attend* (attend left vs. right) could be decoded from both the evoked potentials and alpha power. For decoding based on evoked potentials (i.e., ERP-based decoding), there was strong evidence (BF_10_ > 3) for above chance decoding accuracy from around the time of cue onset. The decoding accuracy increased sharply after stimulus onset and reached a peak at 340 ms after stimulus onset and then went down (but was still above chance until the end of the epoch). There was also strong evidence that information about *where to attend* could be decoded from alpha power starting from the pre-stimulus period (−580 ms), increasing after stimulus onset, peaking at 140 ms, and lasting until the end of the trial.

Information about *what to attend to* (attend colour vs. orientation; Figure 1C) could be decoded from both EEG signals as well, but with different dynamics compared with *where to attend*. The decoding accuracy based on evoked potentials rose gradually following stimulus onset, was robustly above chance from stimulus onset and continued to increase until the end of the epoch. There was also evidence that evoked potentials carried information about *what to attend to* after cue onset for a short period (−1000 to −780 ms). Alpha-based decoding was weaker overall, there was evidence for coding above chance from 660 ms after stimulus onset to the end of the epoch. There was evidence for the null of no alpha-based decoding of *what to attend to* during cue presentation.

For *visual feature information* (Figure 1D), we found evoked potentials coded both attended and unattended features of the attended object, but not features of the unattended object. Decoding began shortly after stimulus onset (∼60 ms) and peaked around 200 ms. By contrast, alpha power did not decode any kind of feature information.

To check the specificity of the alpha decoding results, we conducted exploratory decoding analyses based on power from a broad range of frequencies (5 Hz bands between 13 and 48 Hz). We found at least some evidence that power in other frequency bands, especially low-beta (13-18 Hz), which is adjacent to the alpha frequency range, also carried information about *where to attend* though this decoding tended to be weaker and less sustained than for alpha power, especially in the pre-stimulus period (Figure S1). For comparison with previous literature, we also conducted 1-40 Hz filtered EEG voltage-based decoding. We found highly similar results to ERP-based decoding (Figure S2).

In summary, we found that evoked potentials dynamically coded all the task-relevant information. Information about *where to attend* was present in evoked responses from cue onset. Then at stimulus presentation, information about *where to attend* increased sharply, peaking 340 ms, and was accompanied by coding the *visual features* of the relevant object which peaked slightly before at around 200 ms. Information about *what to attend to* was encoded initially after the cues for a short time (−1000 to −780 ms) and then gradually encoded from around the time of stimulus onset and was strongest at the end of the trial. Alpha power did not code visual feature information, but did carry information about *where to attend* from around the time of the first delay (before stimulus onset) towards the end of the trial and also represented information about *what to attend to* towards the end of the trial (660-1000 ms).

### Attention modulated coding of visual features in an object-based way

To examine how selective attention modulated the representation of visual feature information, we performed timepoint-by-timepoint 2*2 Bayesian repeated measure ANOVAs (factors: object: attended object vs. unattended object; feature: attended feature vs. unattended feature) on ERP-based and alpha-based decoding accuracy of visual feature (Figure 1E). We found that there was evidence for the main effect of object (attended vs unattended) during 160-280 ms (3.06 < BF_10_ < 48.17) for ERP-based decoding, supporting the object-based hypothesis that information deriving from attended objects was encoded more strongly than the features of unattended objects. There was also evidence for a main effect of object in a later period (820-860 ms; 3.06 < BF_10_ < 3.24) but coding of visual features was not above chance at this time. In contrast, for most of the epoch there was evidence towards the null for the main effect of feature or the interaction between object and feature on visual feature information coding, suggesting no prioritisation of attended over unattended features, even within attended objects. The only possible exception was around 600ms, when only stimulus information in attended feature of the attended object was decodable and the Bayes Factors for the interaction were inconclusive. For alpha-based decoding, there was evidence for an interaction between object and feature between 420-580 ms (3.38 < BF_10_ < 6.11), but coding of visual features was not above chance for any condition in alpha power at this time (Figure 1D). There was evidence towards the null for the main effects of object or features on visual feature information coding in alpha power for most of the epoch. Overall these results supported the object-based attention hypothesis that both the attended feature and the unattended feature of the attended object were prioritised.

### Part 2: Concurrent TMS-EEG

Next, we used concurrent TMS-EEG to examine the causal role of right parietal activity in driving these dynamics. We first examined whether the rTMS protocol, compared to the control aTMS, affected alpha power, ITPC, and posterior-contralateral ERP amplitudes, as expected from previous literature. We also checked whether our EEG-only results replicated in the aTMS data. Then, we addressed the key question of whether rTMS affected the evoked and alpha-based decoding of three task aspects (*where to attend, what to attend to,* and *visual features*). Finally, we asked whether the neural changes were reflected in behaviour. For this, we compared the behavioural performance between the two TMS conditions and calculated correlations between TMS-induced decoding changes and TMS-induced behavioural changes.

### Alpha rTMS increased alpha power and ITPC

To check whether our intervention modulated alpha activity, as intended, we examined whether alpha rTMS entrained alpha power and ITPC compared to aTMS using time-frequency decomposition. Compared with aTMS, rTMS showed power and ITPC increase around the right posterior stimulation areas after the first pulse (Figure 2A, 2B).

**Figure 2.**
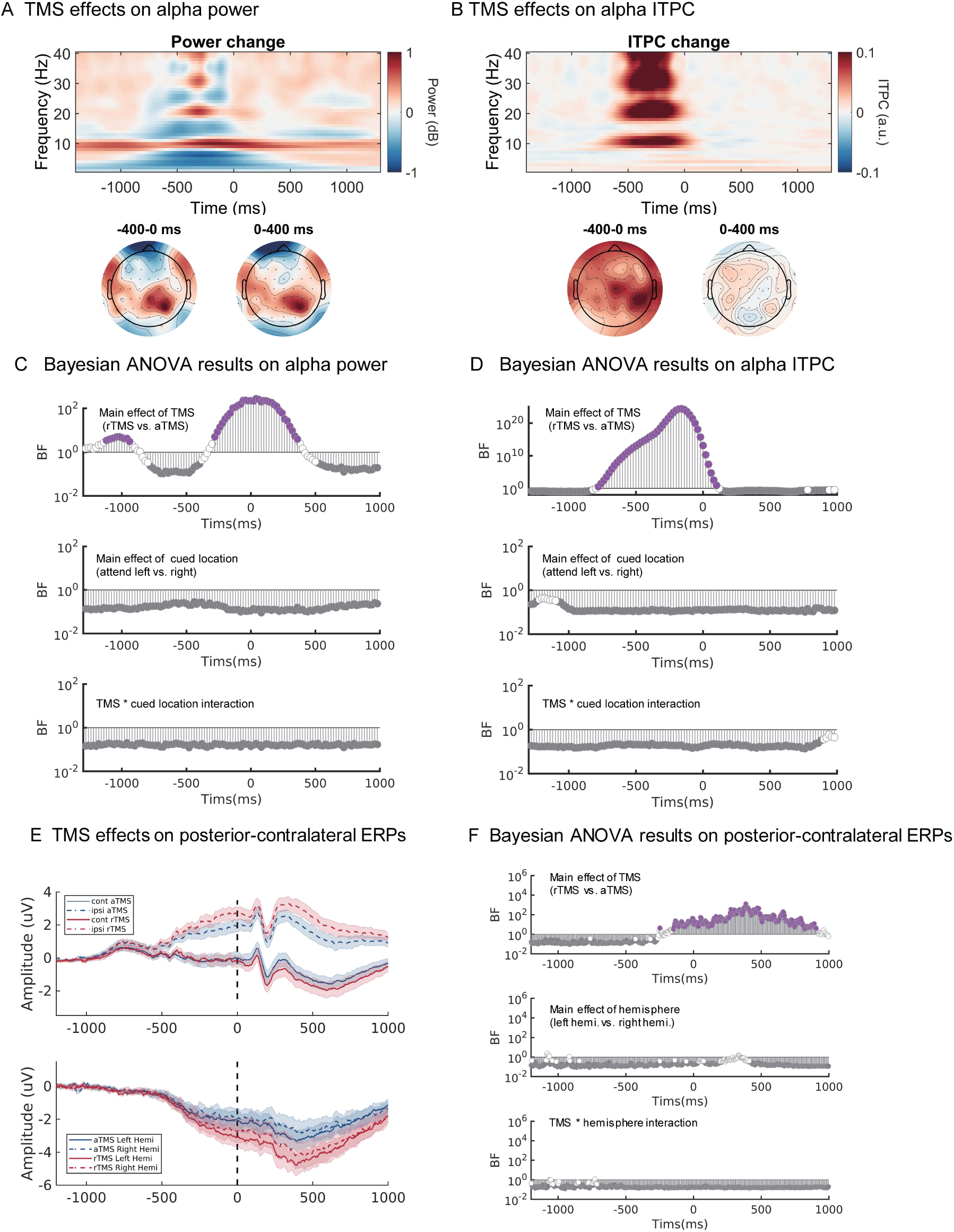
TMS entrained alpha oscillations and affected posterior-contralateral ERP amplitudes. (A) Time-frequency analysis for alpha power entrainment (alpha rhythmic TMS – arrhythmic TMS; averaged across attend left and attend right conditions over right posterior electrodes). The topographic plots show the scalp distribution of the alpha power entrainment effect in time windows before and after stimulus onset. (B) Time-frequency analysis for alpha ITPC entrainment (alpha rhythmic TMS – arrhythmic TMS; averaged across attend left and attend right conditions). The topographic plots show the scalp distribution of the entrainment effect in time windows before and after stimulus onset. (C) Results of repeated-measure Bayesian ANOVAs (TMS: rTMS vs. aTMS; cued location: attended left vs. right) for each timepoint on alpha power. (D) Results of repeated-measure Bayesian ANOVAs (TMS: rTMS vs. aTMS; cued location: attended left vs. right) for each timepoint on alpha ITPC. (E) TMS effects on posterior-contralateral amplitudes. The upper plot shows the evoked responses in the contralateral vs. ipsilateral hemisphere to the spatial cue during different TMS protocols using posterior electrodes (see Methods). The lower plot shows the contralateral minus ipsilateral evoked responses in the left and right hemispheres separately. Baseline correction was performed using the averaged data between -1400 and -1350 ms. Shaded error bars indicate the standard error. (F) Results of repeated-measure Bayesian ANOVAs (factors: TMS: rTMS vs. aTMS; hemisphere: left hemisphere vs. right hemisphere) for each timepoint on evoked potentials (contralateral – ipsilateral wave; the lower plot of Figure 2E). In all plots, time is relative to stimulus onset (ms). Bayes Factor (BF_10_) at each time point was on a log scale, and we marked the BF_10_ with BF_10_ < 1/3 in grey and BF_10_ > 3 in coloured circles. We downsampled the Bayesian ANOVAs results to 200 Hz in for visualisation.

We then performed 2*2 repeated measures Bayesian ANOVAs (TMS: rTMS vs. aTMS; Cued location: Attend left vs. Attend right) on alpha power and ITPC for each timepoint (Figure 2C, 2D). There was strong evidence supporting the main effect of TMS on alpha power from -280 ms before stimulus onset to 360 ms after stimulus onset (3.95 < BF_10_ < 281.84; peaking at around 40 ms after stimulus onset) and on alpha ITPC from -780 ms before stimulus onset to 100 ms after stimulus onset (3.61 < BF_10_ < 2.77×10^24^; peaking at around -160 ms before stimulus onset). This suggested that rTMS affected alpha power both during and beyond TMS pulse period, while it mainly affected ITPC during TMS. There was evidence for the null hypothesis for the main effect of cued location (BF_10_ < 1/3) and interactions between TMS and cued location (BF_10_ < 1/3) on either alpha power or ITPC, indicating that the TMS effects on alpha power and ITPC did not vary with attended location. It should be noted that we found an unexpected main effect of TMS on alpha power during -1120 to -940 ms with medium evidence (3.07 < BF_10_ < 5.06). This might reflect a block-level TMS effect, since the TMS conditions were blocked.

As the rTMS also produced phase-locked neural responses, one possibility was that the change in alpha power reflected transient evoked activity rather than oscillations in alpha band. To exclude this possibility, we subtracted the evoked activity from each trial for each condition (i.e., TMS*Cued location) and performed time-frequency analysis on obtained induced power. As shown in Figure S3A, we still found that alpha power increased, around the target area, under the rTMS condition compared to the aTMS condition. Bayesian ANOVA results similarly showed strong evidence supporting the main effect of TMS on alpha power from -20 ms before stimulus onset to 340 ms after stimulus onset (3.50 < BF10 < 60.65; peaking at around 160 ms after stimulus onset). This result suggested that the rTMS-entrained alpha power (especially after stimulus onset) was induced, rather than just reflecting transient evoked activity.

In sum, this set of analysis showed that our rTMS protocol entrained local alpha power during and after the TMS train, and increased ITPC mainly during the TMS train.

### Alpha rTMS affected posterior-contralateral ERP amplitudes

Posterior-contralateral ERP components (such as N2pc and SPCN) reflect attention allocation to cued locations, and are difference waves calculated by subtracting the waveform from the posterior hemisphere ipsilateral to the cued location from the posterior hemisphere contralateral to the cued location.

To investigate whether rTMS affected posterior-contralateral ERP components, and whether rTMS had a hemisphere-specific effect, we performed 2*2 repeated measures Bayesian ANOVA (with factors TMS: rTMS vs. aTMS; Hemisphere: Left hemisphere vs. Right hemisphere) on amplitudes calculated by subtracting the ERPs at the electrodes contralateral to the cued location from the ipsilateral electrodes, for each timepoint (Figure 2E). The results showed that there was strong evidence of the main effect of TMS on the amplitudes of posterior-contralateral ERP components from around 150 ms before stimulus onset to the end of the trial (−149 to 906 ms; 3.38 < BF_10_ < 1287.95, peaking at 386 ms) (Figure 2F). These time windows cover periods of many classic ERP components like N2pc, SPCN, and anticipatory N2pc-like activities ^55^, which might be univariate contributors to the multivariate decoding results. Amplitudes of these posterior-contralateral ERP components were more negative under the alpha rTMS condition, indicating that participants might allocate more attentional resources in the rTMS condition compared to the aTMS condition. There was evidence supporting the null hypothesis for the main effect of hemisphere and interactions between TMS and hemisphere on ERP amplitudes (BF_10_ < 1/3).

To examine whether rTMS affected posterior-contralateral ERP components during stimulus processing, we examined the ERP results after removing their pre-stimulus differences with a pre-stimulus baseline correction (−100 to 0 ms). There was still strong evidence of a main effect of TMS on the posterior-contralateral ERP amplitudes during time periods covering classic N2pc and SPCN components (271-301 ms, 326-346 ms, 366-416 ms, 446-451 ms, and 466-481 ms; 3.00 < BF_10_ < 63.27, peaking at 386 ms; Figure S4). Amplitudes of these posterior-contralateral ERP components were again more negative under the alpha rTMS condition. Similarly, there was evidence mainly supporting the null hypothesis for the main effect of hemisphere and interactions between TMS and hemisphere on ERP amplitudes (BF_10_ < 1/3).

In summary, we found that alpha rTMS boosted amplitudes of posterior-contralateral ERP components, suggesting that participants may have allocated more attentional resources to the attended location under the rTMS condition.

### Alpha rTMS enhanced neural coding of *where to attend* before and during task performance

Here we examined whether rTMS affected multivariate coding of *where to attend* based either on evoked potentials or alpha oscillations. First, we examined whether the dynamics of representation under our control condition, aTMS, replicated the EEG-only data. Indeed, the ERP-based decoding patterns under aTMS closely replicated the results of EEG-only data (Figure 1B), with strong evidence that information about *where to attend* could be decoded after the cue onset, rising sharply after stimulus onset, and peaking at 360 ms (Figure 3A, blue trace). Similarly, as in the EEG-only data, alpha power coded information about *where to attend* from before stimulus onset (from -220 ms) peaking at 220 ms (Figure 3B, blue trace).

**Figure 3.**
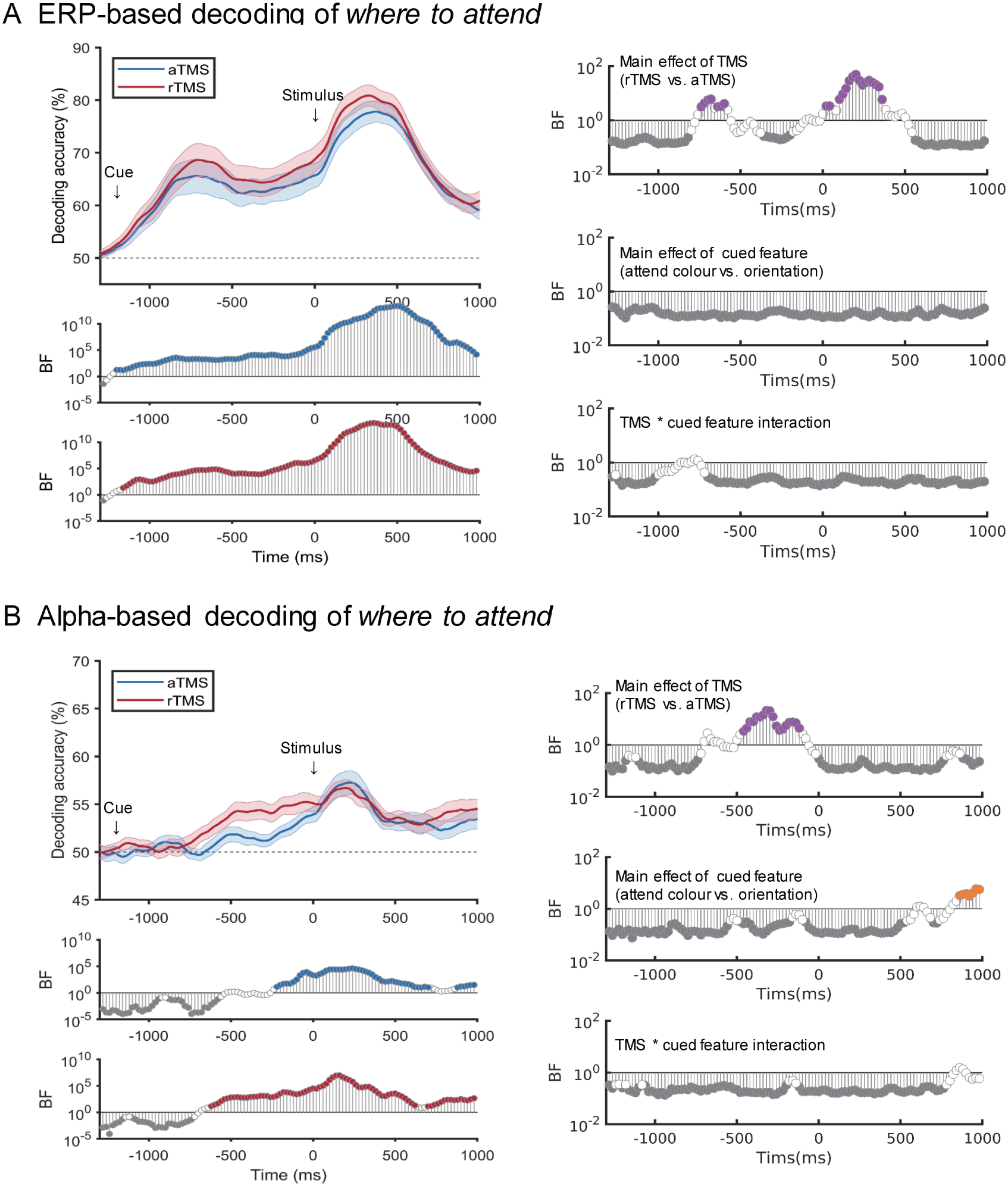
TMS effects on decoding accuracy of *where to attend* (attend left vs. attend right). (A) Left: ERP-based decoding of *where to attend* (attend left vs. attend right) for control aTMS (blue) and alpha rTMS (red). The upper plot shows decoding accuracies, and the middle and lower plots show Bayes Factors (BF_10_) for a comparison of decoding accuracy against chance (50%) at each timepoint, for aTMS (middle, blue) and rTMS (bottom, red). Right: Results of repeated-measure Bayesian ANOVAs (TMS: rTMS vs. aTMS; attended feature: attend colour vs. attend orientation) for each timepoint on decoding accuracy based on evoked potentials. (B) Left: TMS effects on decoding of *where to attend* (attend left vs. attend right) based on alpha power. The upper plot shows decoding accuracies, and the middle and lower plots show Bayes Factors (BF_10_) for a comparison of decoding accuracy against chance (50%) at each timepoint, for aTMS (middle, blue) and rTMS (bottom, red). Right: Results of repeated-measure Bayesian ANOVAs (TMS: rTMS vs. aTMS; attended feature: attend colour vs. attend orientation) for each timepoint on decoding accuracy based on alpha power. Time in all plots is relative to stimulus onset (ms). Cues were shown at approximately -1100 ms (adjusted for IAF). BF_10_ at each time point is shown on a log scale. We marked the BF_10_ with BF_10_ < 1/3 in grey and BF_10_ > 3 in coloured circles.

To quantify the effect of rTMS on coding of *where to attend* (attend left vs. attend right), we performed 2*2 repeated-measure Bayesian ANOVAs (factors: TMS: rTMS vs. aTMS; attended feature: attend colour vs. attend orientation) for each timepoint and each decoding method (Figure 3). As can be seen in Figure 3A, ERP-based decoding accuracy tended to be higher for rTMS (red) compared to aTMS (blue) from around the time of TMS onset lasting through and beyond stimulus onset. The ANOVA revealed medium evidence for the main effect of TMS around the onset of the TMS pulses (∼ -740 to -600 ms from stimulus onset; 3.17 < BF_10_ < 6.10), and medium to very strong evidence for an effect of TMS after stimulus onset (∼ 100 to 360 ms from stimulus onset; 5.96 < BF_10_ < 50.14, peaking at 200 ms). The was evidence for the null for the main effect of cued feature and the interaction between TMS and cued feature, indicating that the cued feature did not influence the information coding about *where to attend* or the effect of TMS on this information.

rTMS also enhanced alpha-based decoding of *where to attend*, but only during the pre-stimulus period (Figure 3B). ANOVA results showed that there was medium to strong evidence for the main effect of TMS before stimulus onset (−460 to −120 ms from stimulus onset; 3.40 < BF_10_ < 22.19; peaking at -300 ms). Also, there was medium evidence supporting the main effect of cued feature at the late period (860 to 1000 ms from stimulus onset, 3.26 < BF_10_ < 5.63), indicating decoding of *where to attend* was higher for the attend orientation condition than the attend colour condition at this time. There was no compelling evidence for the interaction effect between TMS and cued feature on *where to attend* in alpha-power based decoding.

To exclude the possibility that alpha-based decoding results might originate from the contributions of pulse-evoked, non-oscillatory components, we also performed alpha-based decoding analysis using the data after subtracting evoked activity from each trial for each condition. As shown in Figure S5, the decoding results and the ANOVA results were both similar to the results shown in Figure 3B, suggesting that the decoding results of alpha power were from induced alpha oscillations rather than evoked activities.

### Alpha rTMS did not influence neural coding of *what to attend to* or task-relevant *visual feature information* during task performance

For information about *what to attend to*, the decoding patterns under our control aTMS condition (Figure 4, blue trace) were again similar to the pattern of EEG-only data (Figure 1C). Specifically, ERP-based decoding accuracy of *what to attend to* became robustly above chance from around the time of stimulus onset and lasted until the end of the epoch (Figure 4A, blue trace). Alpha power, in contrast, coded this information at the late time period after around 700 ms from stimulus onset (Figure 4B, blue trace). These results indicated the patterns reported above in the EEG-only data were replicable, and that the aTMS condition acts as a suitable control for our rTMS condition of interest.

**Figure 4.**
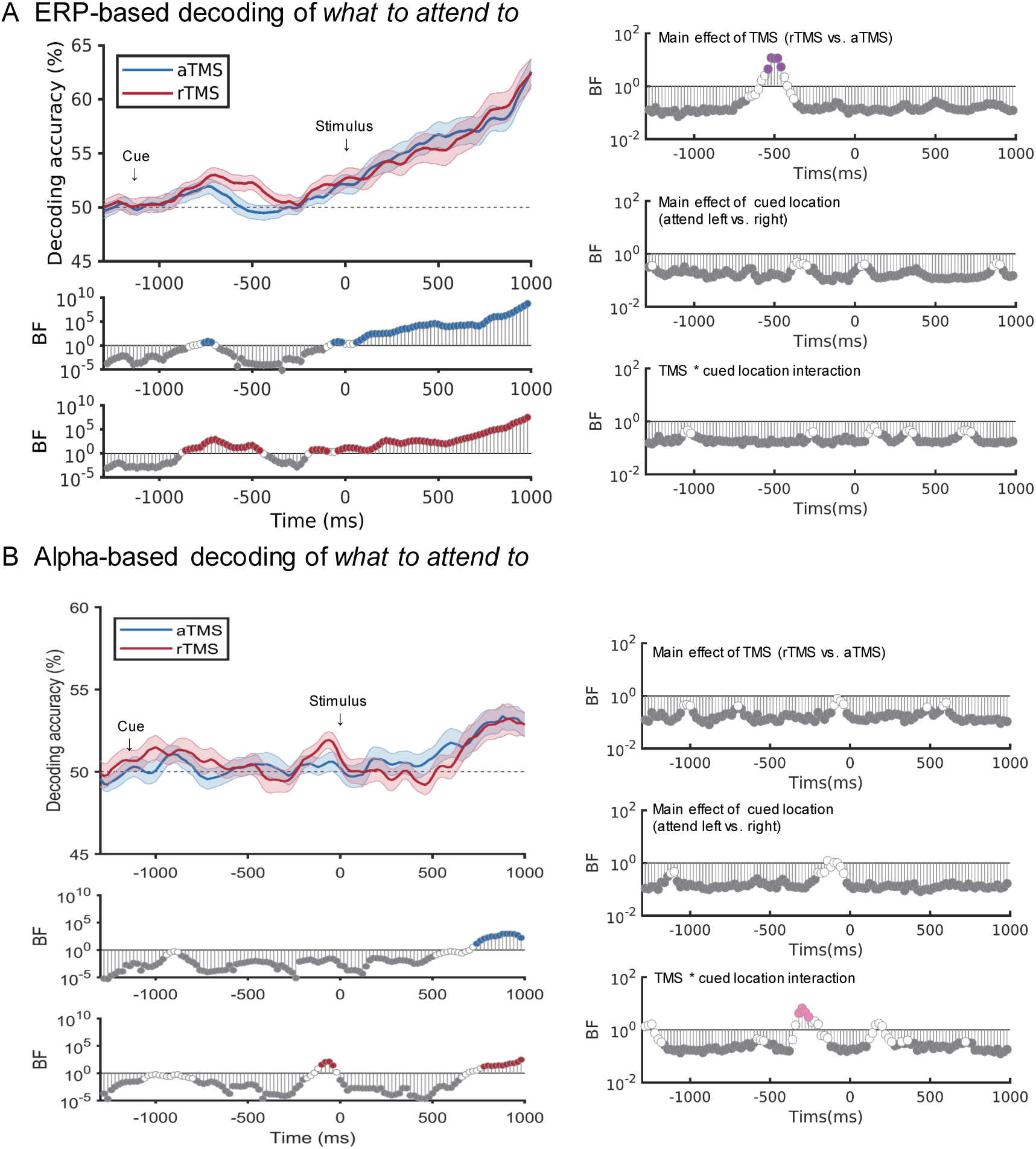
TMS effects on decoding accuracy of *what to attend to*. (A) Left: ERP-based decoding of *what to attend to* (attend colour vs. attend orientation) for control aTMS (blue) and alpha rTMS (red). The upper plot shows decoding accuracies, and the middle and lower plots show Bayes Factors (BF_10_) for a comparison of decoding accuracy against chance (50%) at each timepoint, for aTMS (middle, blue) and rTMS (bottom, red). Right: Results of repeated-measure Bayesian ANOVAs (TMS: rTMS vs. aTMS; attended location: attend left vs. attend right) for each timepoint on decoding accuracy based on evoked potentials. (B) Left: Alpha power-based decoding of *what to attend to* (attend colour vs. attend orientation) for control aTMS (blue) and alpha rTMS (red). The upper plot shows decoding accuracies, and the middle and lower plots show Bayes Factors (BF_10_) for a comparison of decoding accuracy against chance (50%) at each timepoint, for aTMS (middle, blue) and rTMS (bottom, red). Right: Results of repeated-measure Bayesian ANOVAs (TMS: rTMS vs. aTMS; attended location: attend left vs. attend right) for each timepoint on decoding accuracy based on alpha power. Time in all plots is relative to stimulus onset (ms) and the cues were shown at approximately -1100 ms. BF_10_ at each time point is marked on a log scale. We marked the BF_10_ for these results with BF_10_ < 1/3 in grey and BF_10_ > 3 in coloured circles.

To quantify the effect of rTMS on information coding about *what to attend to* (attend colour vs attend orientation), we conducted 2*2 repeated-measure Bayesian ANOVAs (TMS: rTMS vs. aTMS; cued location: attend left vs. attend right) for each timepoint and each decoding method. ANOVA results for ERP-based decoding (Figure 4A) and alpha-based decoding (Figure 4B), both largely showed evidence supporting the null hypothesis for either main or interaction effects. The exception was medium evidence for the main effect of TMS on ERP-based decoding during TMS pulses (−540 to −460 ms from stimulus onset; 4.52 < BF_10_ < 11.98), showing decoding accuracy for *what to attend to* was higher in the rTMS condition than that in the aTMS condition. However, this effect was short and did not endure past stimulus presentation into the period of task performance. Also, there was a short time window with evidence supporting interaction effect on alpha-based decoding before stimulus onset (−320 to −260 ms from stimulus onset; 3.15 < BF_10_ < 6.70). This interaction suggested that there was a larger effect of TMS on decoding of *what to attend to* when participants attended left, but post-hoc Bayesian t-tests did not provide evidence for TMS differences for either attend left (1.72 < BF_10_ < 2.87) or attend right conditions (0.40 < BF_10_ < 0.45) in this time window.

In summary, there was a brief effect of TMS during TMS presentation, indicating that decoding of *what to attend to* was more sustained during the rTMS train than during the aTMS train. However, during task performance (after stimulus presentation) there was evidence for the null hypothesis that TMS did not affect coding of *what to attend to*.

For coding of *visual feature information* (Figure 5A, 5B), the decoding patterns under aTMS condition (Figure 5, blue traces) again replicated the EEG-only data. Only evoked potentials, rather than alpha power, coded feature information of the attended object after stimulus onset lasting around 200 ms.

**Figure 5.**
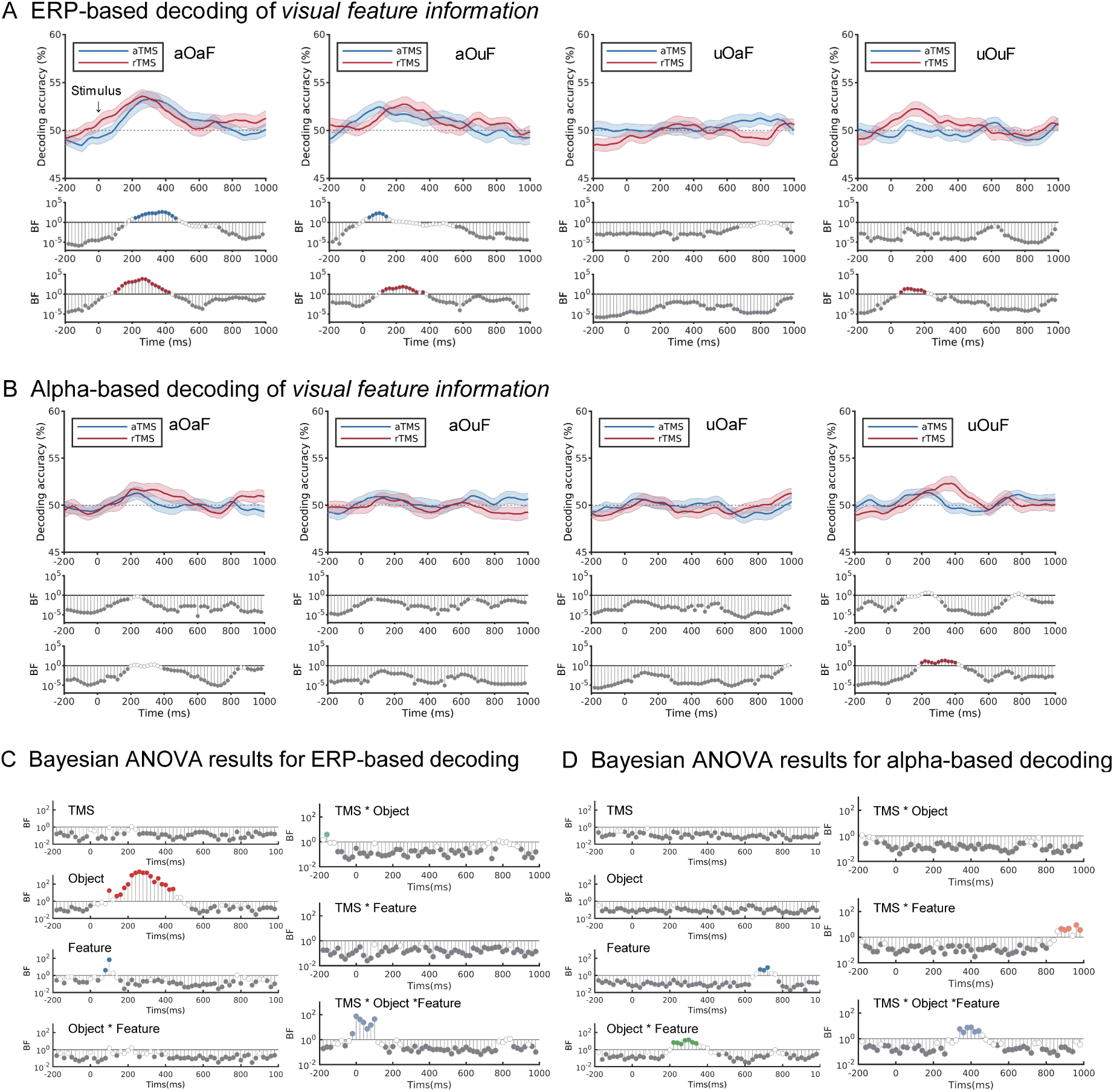
TMS effects on decoding accuracy of *visual feature information*. (A) ERP-based decoding of *visual feature information* for control aTMS (blue) and alpha rTMS (red), in each attention condition separately. Upper plots show decoding accuracy and middle and lower plots show the associated Bayes Factors (BF_10_) for a comparison of decoding accuracy against chance (50%) at each timepoint. (B) Alpha power-based decoding of *visual feature information* for control aTMS (blue) and alpha rTMS (red), in each attention condition separately. Upper plots show decoding accuracy and middle and lower plots show the associated Bayes Factors (BF_10_) for a comparison of decoding accuracy against chance (50%) at each timepoint. (C) Results of repeated-measure Bayesian ANOVAs (TMS: rTMS vs. aTMS; object: attended object vs. unattended object; feature: attended feature vs. unattended feature) for each timepoint on decoding accuracy based on evoked potentials. (D) Results of repeated-measure Bayesian ANOVAs (TMS: rTMS vs. aTMS; object: attended object vs. unattended object; feature: attended feature vs. unattended feature) for each timepoint on decoding accuracy based on alpha power. aOaF: attended feature of attended object; aOuF: attended feature of unattended object; uOaF: unattended feature of attended object; uOuF: unattended feature of attended object. Time in all plots is relative to stimulus onset (ms). BF_10_ is shown on a log scale. We marked the BF_10_ for these results with BF_10_ < 1/3 in grey and BF_10_ > 3 in coloured circles.

To quantify the effect of rTMS on information coding about *visual feature information* and to examine any interaction between rTMS and how attention affects visual information coding, we conducted 2*2*2 repeated-measure Bayesian ANOVAs (factors: TMS: rTMS vs. aTMS; object: attended object vs. unattended object; feature: attended feature vs. unattended feature) for each timepoint and each decoding method. There was evidence for the null hypothesis regarding the main effect of TMS for both ERP-based and alpha-based decoding at almost all timepoints (Figure 5C, 5D). For ERP-based decoding, TMS also did not interact with the effect of object or attention. There was a 3-way interaction during -20 to 100 ms, reflecting a larger object*feature interaction under rTMS compared to aTMS. However, the evidence for an object*feature interaction was weak in both TMS conditions (all BF_10_ < 4.14). For alpha-based decoding there was evidence for an interaction between TMS and feature, but only at late timepoints where coding was not above chance in any condition. There was also a 3-way interaction during 340 to 440 ms, reflecting a larger object*feature interaction under rTMS compared to aTMS. This effect was likely driven by above chance coding in the uOuF rTMS condition during 200 to 400 ms. However, the overlapping period between this evidence and the above chance uOuF coding lasted only for 4 timepoints, so we decline to draw any strong inference from it. In summary, we did not find meaningful effect of rTMS on coding of task-relevant *visual feature* information.

Similar to EEG-only data, we found strong evidence supporting the main effect of object on ERP-based decoding accuracy during the period 100 to 440 ms after stimulus onset (3.96 < BF_10_ < 2715.19, Figure 5C, red) and no compelling evidence for either the main effect of feature or interaction between object and feature. There was evidence towards a main effect of feature around 100 ms after stimulus presentation (Figure 5C, dark blue), but this evidence lasted for only for 2 timepoints. In the alpha-based decoding there was evidence for a feature*object interaction at 220-340 ms (4.53 < BF_10_ < 13.49, Figure 5D, green) and weak evidence for a main effect of feature (3.80 < BF_10_ < 8.32, Figure 5D blue) lasting for 3 consecutive timepoints in the alpha-based decoding, but we do not think these effects were meaningful as the decoding based on alpha power was generally not above chance. These results, especially for aTMS, are consistent with the EEG-only data, supporting the object-based attention hypothesis that attention enhances processing of both relevant and irrelevant features of attended objects.

To sum up, there was evidence supporting the null hypothesis that alpha rTMS over IPS did not affect coding of what participants should attend to or coding of the associated visual feature information, particularly during the period of task processing (after stimulus onset). This suggests that the causal effect of pre-stimulus parietal alpha rTMS, on alpha power-based and ERP-based coding, during task performance, was specific to information about *where to attend*.

### Behavioural effects of rTMS

The behavioural results for both TMS conditions are shown in Table S1. To examine whether rTMS had an overall effect on participants’ behavioural performance on the selective attention task, we first performed 2*2*2 repeated measure Bayesian ANOVAs (factors: TMS: rTMS vs. aTMS; cued location: attend left vs. attend right; cued feature: attend colour vs. attend orientation) on participants’ RT and response error data. As in the EEG-only data, there was strong evidence for a main effect of cued feature on participants’ RT and response error, which reflect the two different tasks and response methods. Participants had higher response error (BF_10_ = 7.77×10^15^) but shorter RT (BF_10_ = 4.82×10^28^) in the colour task compared to the orientation task. Other than the main effects of cued feature, all other effects showed evidence for the null (all BF_10_ < 1/3).

Even in the absence of an overall (group-level) effect, it is possible to test whether between-subject variation in response to TMS is meaningful. In particular, given the specific role of rTMS on modulating coding of *where to attend*, we were interested to know whether the rTMS-induced change in decoding was reflected in behavioural performance. To examine this, we performed timepoint-by-timepoint rank correlations (Kendall’s Tau) between TMS-induced decoding changes on *where to attend* and the TMS-induced changes in response error and RT, and then calculated the one-sided Bayes Factor for the correlation coefficients ^56^ (Figure 6). The results showed that, at several points in the trial, TMS-induced changes in both ERP-based and alpha-based decoding of *where to attend* negatively correlate with the TMS-induced changes in response error and RT. The direction of the correlation indicated that increases in decoding correlated with better behavioural performance (lower response error or shorter RT). Specifically, TMS-induced improvement in response accuracy was associated with increased ERP-based decoding in a short period before stimulus onset (−320 to −260 ms after stimulus onset; 3.13 < BF_-0_ < 6.26) and in the time period from 480 to 680 ms after stimulus onset (3.13 < BF_-0_ < 10.72, Figure 6A, pink), and with increased alpha-based decoding from 180-280 ms (3.13 < BF_-0_ < 8.47, Figure 6B, pink). TMS-induced improvements in RT were associated with TMS-induced improvement in ERP-based decoding, most strongly around 80-300 ms after stimulus onset (3.13 < BF_-0_ < 13.74, Figure 6A, green), and with TMS-induced improvement in alpha-based decoding towards the end of the epoch (840-1000 ms; 3.13 < BF_-0_ < 6.26, Figure 6B, green). These findings suggest that while rTMS did not produce an overall effect on behavioural performance, it did influence response error and RT at the individual level, in a manner that can be predicted from its influence on coding of *where to attend*. It also highlights the association between ERP- and alpha-based decoding and behaviour.

**Figure 6.**
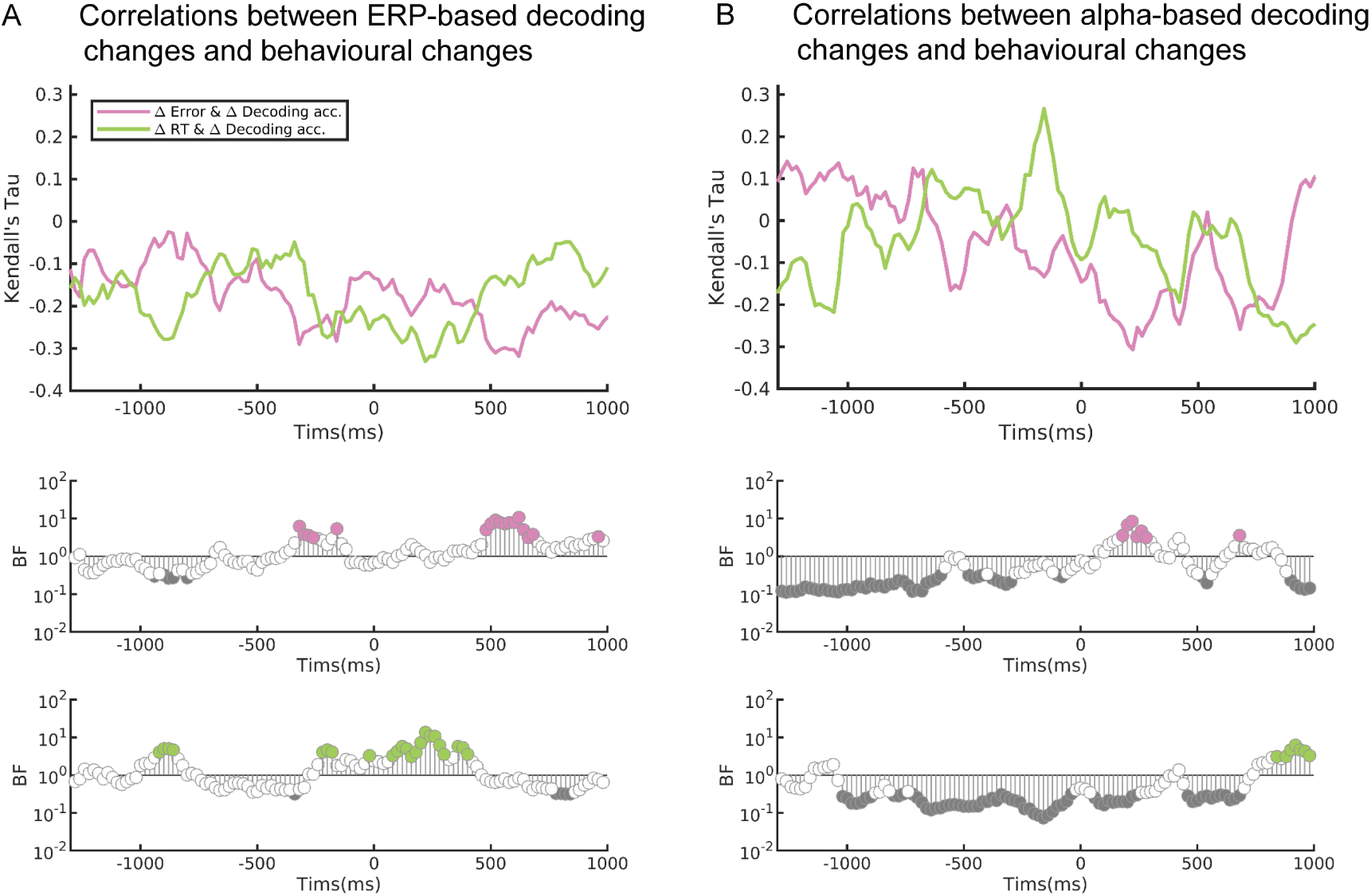
Correlations (Kendall’s Tau) between TMS-induced changes in decoding of *where to attend* and TMS-induced changes in behavioural response. (A) Correlations between TMS-induced ERP-based decoding changes of *where to attend* and response error (pink) and RT (green). Negative values of Kendall’s Tau reflect an association between a TMS-induced increase in decoding and with TMS-induced improvement in behavioural performance (lower response error or shorter RT). TMS-induced improvement in error rate was associated with TMS-induced improvement in decoding shortly before stimulus presentation and around 500ms after stimulus onset. TMS-induced improvement in reaction time was associated with TMS-induced improvement in decoding for around 300ms after stimulus onset. (B) Correlations between TMS-induced alpha-based decoding changes of *where to attend* and response error (pink) and RT (green). Negative values of Tau reflect an association between a TMS-induced increase in decoding and with TMS-induced improvement in performance (lower response error or shorter RT). TMS-induced improvement in error rate was associated with TMS-induced improvement in alpha-based decoding around 200ms after stimulus onset. TMS-induced improvement in reaction time was associated with TMS-induced improvement in decoding for around 840ms after stimulus onset. One-sided Bayes Factor (BF_-0_) at each time point is plotted on a log scale. We report one-sided tests as we pre-specified the direction that larger decoding improvement should predict better performance. We marked the BF_-0_ < 1/3 in grey and BF_-0_ > 3 in coloured circles.

## Discussion

This study aimed to explore the dynamics with which three different aspects of a selective attention task (*where to attend*, *what to attend to*, and *visual feature information*) are coded in evoked potentials and alpha oscillations, and the causal role of right parietal cortex in supporting these representations. We first employed multivariate decoding methods that used alpha power and evoked potentials to investigate their respective contributions to selective attention processing. Then, we applied a TMS entrainment protocol, with concurrent EEG recording, to manipulate activity in the right IPS and read out the effects on neural activity. Results showed that evoked potentials coded all aspects of task-relevant information at different time points while alpha power only coded information about *where to attend* and *what to attend to*. When we turned to causal intervention, we found that our alpha rTMS protocol entrained local alpha power and ITPC, boosted posterior-contralateral ERP amplitudes, and more importantly, enhanced the neural coding of information about *where to attend*. Furthermore, the TMS-induced decoding changes in *where to attend* predicted the changes in behavioural RT and response error. These findings provide insights into the function of evoked potentials and alpha oscillations in attentional selection and suggest that IPS-controlled evoked potentials and alpha oscillations play a causal and specific role in supporting coding of information about *where to attend* during selective attention.

Both EEG-only and TMS-EEG results demonstrated that evoked potentials represent all the three types of task information with a dynamic representation scheme. Information about *where to attend* was coded after cue onset and maintained towards the stimulus onset. Following the stimulus onset, coding of information about *where to attend* increased further, and was accompanied by information about *visual feature* (which peaked first) and *what to attend to* (which peaked at the end of the trial). The data add to a growing body of literature suggesting dynamic coding schemes where multiple task elements must be computed and/or integrated ^26,28^. The specific order and dynamics with which various task elements are represented seems likely to vary depending on the details of the task, and further work will be needed to draw out any common principals, but dynamic schemes seem attractive for avoiding interference when multiple task aspects must be processed and integrated.

Decoding based on evoked potentials may pick up on neural signals that are also reflected in traditional univariate ERP components. For example, the N2pc is known to be a marker of attentional allocation to specific cued locations ^12,57^, and larger negative responses in the hemisphere contralateral to the attended location would provide a signal that a multivariate classifier could use to determine where attention was being allocated. Of course, ERP-decoding for information about *where to attend* showed a longer time window than any specific ERP components, suggesting that decoding of *where to attend* is sensitive to many elements of the evoked neural response. Compared to univariate ERPs, decoding has increased sensitivity to detect differences between conditions ^25,49^, although this sensitivity comes at the price of reduced specificity about what univariate changes the decoding reflects, particularly as it drops the requirement for the change to be in the same direction across participants ^58^. This means that decoding may reflect more complex patterns of activation change than is typically visible when considering the group response at each channel separately. More work could be done to understand the relationship between multivariate decoding and traditional univariate ERP components in different situations.

Alpha power also showed a dynamic coding scheme, with early coding of *where to attend* (from pre-stimulus period) and later coding of *what to attend to* (at the end of the epoch). The observation of alpha-based coding of *where to attend* echoes previous studies showing that alpha power robustly reflects spatial location information regarding *where to attend* ^20,59^. It is also consistent with the idea of the spatial spotlight theory ^21,60,61^ that alpha indexes the spatial allocation of attention across the visual field. However, the observation that alpha power also carried information about *what to attend to* during the later time period was novel, and suggests that alpha power may have the potential to carry different types of information at different processing stages. Meanwhile, in our data alpha power could not decode feature information regardless of whether it was task-relevant or irrelevant. This is consistent with previous findings in working memory tasks that both evoked ERPs and alpha oscillations code spatial information, while only evoked potentials also code visual feature information^8,60,62^.

When we examined how attentional states modulated the coding of visual feature information, both EEG-only and TMS-EEG data showed that the evoked potentials code both attended and unattended visual features of the attended object. This result is consistent with object-based attention theories ^32–34^ which predict that attention spreads to support processing of multiple features of attended objects. This result is, however, not consistent with our previous MEG decoding results suggesting a multiplicative coding pattern in which only the attended feature of the attended object is attended ^26,37^ or studies reporting a benefit for task-relevant features across the visual field ^6,7,36^. A possible basis for this discrepancy could be different perceptual load or complexity of the stimuli between studies. In our studies supporting a multiplicative account, the perceptual load of the stimuli was high (see ^26^), which may drive more selective processing of only the most relevant features. In contrast, the stimuli in the present study had a much lower perceptual load, so participants might have more spare resources deployed to the other feature of the attended object ^63,64^. Indeed, we have previously suggested that participants may not prioritise neural processing of task-relevant features of single objects when task demands are low ^37^. Additionally, in the current task the potential for interference from the irrelevant dimension was lower than in the designs where we have observed multiplicative effects. In essence, we suggest that both (or even all three) accounts may fit in different circumstances, and allocation of attentional resources to features and objects might be flexible to match the requirements of the current task.

Relative to an arrhythmic control, rTMS at IAF over right parietal cortex entrained local alpha power and ITPC as expected, consistent with previous TMS entrainment studies ^43,45^. It also induced more negative amplitudes for posterior-contralateral ERP components, perhaps indicating that participants allocated more attentional resources to the attended location under the rTMS condition ^12,18,55^. This was also suggested by the rTMS-induced increase in decoding of *where to attend* based on evoked potentials and alpha power, which in turn predicted the degree of individual-subject improvement in behavioural performance. These findings underscore the causal role of parietal alpha in supporting the allocation of selective attention in space.

Our data illuminate the dynamics with which entraining pre-stimulus parietal alpha affects coding of *where to attend*. The supportive effect of rTMS on decoding started during rTMS presentation in the preparatory period and was sustained through task performance. It boosted first ERP-based and then alpha-based decoding during the preparatory period, and then boosted ERP-based decoding through the first few hundred milliseconds of task performance (after stimulus presentation). Previous studies and theory have suggested that posterior alpha power is sensitive to spatial cues ^21,60,65^, and when participants’ attention is allocated to different locations, the scalp distribution of posterior alpha systematically shifts ^8,59^. This can be elicited by spatial cues and is commonly known as anticipatory alpha ^21,66^, which may be related to our finding that rTMS increased alpha-based decoding of *where to attend* before stimulus onset (−460 to −120 ms from stimulus onset). After stimulus presentation, however, the effect of rTMS was translated into ERP-based decoding (∼100 to 360 ms after stimulus onset). This may reflect similar mechanisms as reflected in TMS-based modulation of the posterior-contralateral ERP amplitudes like the N2pc ^46^ and potential relationship between alpha and these posterior-contralateral ERPs ^66^. More studies are needed to explore the mechanisms by which pre-stimulus alpha rTMS induces ERP changes.

We did not find compelling alpha rTMS effects on the neural coding of information about *what to attend to* or *visual features* except a short boost for *what to attend to* shortly after the first few TMS pulses (∼-540 ms to -460 ms) which did not endure into the time period of task performance. Our interpretation is that the pre-stimulus alpha oscillations at right IPS play a specific role in supporting the allocation of attention in space, which aligns with previous studies on alpha and IPS functions in selective attention ^13,67^. However, we cannot rule out the possibility that TMS at another point in the trial (such as late in the trial, when alpha-power coded for *what to attend to*) would affect coding of other task features. In future studies, it would be interesting to investigate whether different timing of alpha rTMS can produce distinct effects on neural coding and behaviour, and whether oscillations and or evoked responses in other brain areas can be modulated to enhance the representation of other task features.

Similar to many previous TMS studies ^68,69^, we did not observe TMS effects on overall behavioural performance, potentially reflecting the known large individual differences in response to TMS protocols ^70^. However, we did find that TMS-induced decoding changes (for both ERP- and alpha-based) in information about *where to attend* predicted individual-participant TMS-induced behavioural error changes (both response accuracy and RT). These results add confidence that the TMS-induced changes in coding of *where to attend* were not epiphenomenal, but were in fact behaviourally-meaningful and supported task performance.

In conclusion, our results reveal the dynamics of evoked potentials and alpha oscillations in coding multiple aspects of a selective attention task (i.e., *where to attend*, *what to attend to*, and *visual feature information*), and how they are modulated by posterior pre-stimulus alpha. We found a dynamic coding scheme in which evoked potentials represented all aspects of the task set (*where to attend* and *what to attend to)* as well as the features of attended visual objects, while alpha power carried information about *where to attend* and *what to attend to*. Using concurrent TMS-EEG, we found that right parietal alpha rTMS entrained alpha power and ITPC and boosted posterior-contralateral evoked potentials. Moreover, pre-stimulus alpha rTMS specifically and causally enhanced neural codes for *where to attend* in both evoked activity and alpha power during stimulus processing, and which in turn predicted behavioural performance. This suggests that information about multiple task aspects is carried in evoked and alpha-oscillatory activity, and that IPS-controlled alpha and evoked potentials are specifically important to neural coding of *where to attend*, in service of successful behaviour in selective attention tasks.

## Methods Participants

We recruited 35 healthy right-handed participants with normal or corrected to normal vision from the local community and the online participant database (SONA) at the University of Cambridge. All participants gave written informed consent to participate in the study, which was approved by the by the Cambridge Human Biology Research Ethics Committee. We screened all participants for contraindications for TMS and MRI, including history of epilepsy and/or fainting, current medications, and recent alcohol or drug consumption. Following visual inspection of the raw data, three participants were excluded from TMS-EEG analysis due to excessive motion artefacts in EEG data, and an additional participant was excluded from the EEG-only analysis due to excessive movement during EEG-only data collection. As a result, 32 participants (19-36 years, 18 females, 14 males) were included in TMS-EEG analysis, and 31 of these (19-36 years, 18 females, 13 males) were included in EEG-only data analysis.

## Task design

We used the selective attention task shown in Figure 1A. On each trial, two peripheral tilted coloured bars were presented on the screen, with one bar placed to the left and the other to the right of the central fixation cross. Bars were centred at an eccentricity of approximately 7.8° visual angle and were 4.2° in length and 0.4° in width. The colour of each bar was either yellowish (RGB ranging from 230, 176, 69 to 179, 230, 69; forty degrees across the colour wheel) or blueish (RGB ranging from 69, 176, 230 to 69, 72, 230; forty degrees across the colour wheel). The orientation of each bar was either leftward (varying between 26 and 65 degrees) or rightward (varying between -65 and -26 degrees). Thus, there were 16 combinations of stimuli crossing two categories of colour and two categories of orientation at each of the two locations.

Each trial began with the white fixation cross (randomly displayed for 500-1000 ms) on a grey screen (RGB: 100, 100, 100), followed by a cue presented in the centre of the screen (500 ms). The cue consisted of two symbols that instructed both the location that participants should attend to (‘<’ meant attend left; ‘>’ meant attend right) and the feature that participants should attend to (‘c’ meant attend colour; ‘o’ meant attend orientation). In TMS-EEG blocks, TMS pulses were given during the first delay after the cue. The duration of the first delay varied depending on the participant’s individual alpha frequency (IAF), being either (100+5000/IAF) ms or (100+6000/IAF) ms to align stimulus onset to the first or the second cycle peak after the pulses. For example, if a participant’s IAF was 10 Hz, the duration of the first delay was either 600 ms or 700 ms. Two durations were used (50% of trials each) to avoid a precise temporal prediction of when the stimulus would occur. After the first delay, the stimuli were presented on each side of the screen for 100 ms. Then, there was a second delay of 1000 ms, after which participants saw a response screen and made their responses. In the colour task, participants used a mouse to select the colour of the cued target object in a colour wheel within 5 seconds. The orientation of the colour wheel varied over trials, to make sure that participants could not prepare a motor response. In the orientation task, participants needed to manipulate a black tilted bar at the centre of the screen to match the orientation of the cued target within 5 seconds. The initial orientation of the tilted bar also varied over trials to avoid response preparation. Thus, although the stimuli were later grouped into categories of yellowish/blueish and leftward/rightward for the purposes of our analyses, the task required participants to report precise colour and orientation. The inter-trial interval varied based on response time, so as to fix the sum of the two at 5 seconds. Participants were instructed to fixate on the cross at the centre throughout the experiment, except during the response period.

### Procedure

Participants first attended one MRI session to acquire a structural scan for neuronavigation purposes, unless a pre-existing structural scan was available. They then completed two TMS-EEG sessions, with 5 to 20 days separating the two. Each session was for one TMS condition (alpha rTMS or aTMS) with the order counterbalanced over participants. During each session, participants first practised the task for ∼5 minutes and received feedback on each trial. After practice, we measured participants’ motor threshold and prepared the EEG setup. Then, participants performed three blocks of 64 trials with EEG recording (but no TMS), after which they had a break while we analysed the EEG data to obtain their IAF and used neuronavigation to locate the TMS target. Participants then completed five blocks of TMS-EEG trials, with each block incorporating 64 trials, encompassing all possible combinations of 16 stimuli, 2 cued locations, and 2 cued features. This resulted in the completion of 192 EEG-only trials and 320 TMS-EEG trials per session, for a total of 384 EEG-only trials (used for EEG-only analysis in the first part of the results) and 640 TMS-EEG trials (used for TMS-EEG analysis in the second part of the results).

### MRI acquisition

T1-weighted MRI structural scans were acquired using a Siemens 3T Prisma scanner at the MRC Cognition and Brain Sciences Unit, University of Cambridge. A standard 32-channel head coil was used to acquire MPRAGE structural scans (acquisition time = 4.32 minutes, TR = 2.25 s, TE = 900 ms, flip angle = 9 degrees, FOV = 256 mm × 240 mm × 192 mm, resolution = 1 mm isotropic, acceleration factor = 2).

### TMS protocols

Participants’ motor threshold was measured before the start of the first TMS-EEG session. The resting motor threshold of the relaxed contralateral abductor pollicis brevis muscle was defined as the lowest stimulation intensity capable of causing a visible twitch in the muscle on five out of ten pulses ^71^. The stimulation site for the right IPS was at the Montreal Neurological Institute (MNI) coordinates (*x* = 37, *y* = -56, *z* = 41) derived from the peak activity coordinates of the IPS defined in the multiple-demand control network from previous fMRI studies ^72–74^. The MNI coordinates were transformed into native space for each participant based on their structural scan, which we normalised using the segment and normalise routines of SPM12. During the TMS-EEG sessions, we performed neuronavigation using a Brainsight 2 system to ensure accurate targeting of the stimulation site on each participant by co-registering the head to the structural scan. TMS pulses were delivered using a DuoMag XT-100 stimulator and a figure-of-eight butterfly coil (DuoMag 70BF).

As shown in Figure 1A, participants received a train of five alpha rTMS or aTMS pulses at 110% of their motor threshold (63% of the maximal stimulator output on average), beginning 100 ms after the first delay onset on each trial. For the alpha rTMS condition, the frequency of pulses was the IAF of the specific participant. As a control condition, aTMS had the same total duration as the rTMS for each participant by fixing the timing of the first and the fifth pulse. Each arrhythmic train was randomly generated with each pulse separated by at least 20 ms. In generating these control trains, we checked each train for its proximity to alpha frequency pulse patterns and discarded pulse patterns where all four intervals had values between 80-130 ms and replaced this train with a new pattern. In total, there were 1600 pulses in each TMS-EEG session (either rTMS or aTMS), with an inter-train interval of at least 7.2 seconds. This protocol complies with safety recommendations for TMS protocols ^75^.

### Behavioural data analysis

We quantified behavioural performance as the response error between the cued object’s orientation (or colour) and the reported orientation (or colour) and the time taken to give this response (RT). For the colour task, the error range is from -180° to 180°; for the orientation task, the error ranges from -90° to 90°. Trials on which participants did not respond within the 5 second time limit were discarded from further analyses.

### EEG recording and preprocessing

We used TMS-compatible EEG equipment with (actiCHamp Plus 64 System with actiCAP slim electrode cap 64-channels; Brain Products UK) to continuously acquire EEG data from 64 active electrodes at the scalp (25000 Hz sample rate for TMS-EEG; 1000 Hz sample rate for EEG-only). Channel Fz was used as an online reference. We maintained skin/electrode impedance below 20 kΩ, in line with the manufacturer’s recommendation. Participants wore earplugs to reduce auditory contamination of EEG by TMS clicks and to comply with the safety requirement for hearing protection. We preprocessed EEG data using the EEGLAB toolbox ^76^ with an open-resource extension, TMS-EEG signal analyser (TESA) ^77^, and custom-made Matlab code.

For EEG-only data, we cut the data into 2.9 second epochs (−1.6 to +1.3 seconds from stimulus onset) and applied baseline correction using data from the fixation period (−1.6 to −1.2 seconds from stimulus onset). Then, we filtered the data with a band-pass (0.01-100 Hz) and a stop-band filter (48-52 Hz) using the Butterworth method. Bad trials were visually identified and removed, and bad channels were detected on an epoch-by-epoch basis using the TBT toolbox ^78^. We rejected channels that had more than 30% of bad trials and interpolated them at the end of the preprocessing steps. Next, we conducted independent component analysis (ICA) to identify and remove artefacts reflecting eye movement, blink-related activity, electrode artefacts. Specifically, we used a fastICA algorithm (*pop_tesa_fastica* function) and automatic component classification (*pop_tesa_compselect* function). Each component underwent manual inspection and reclassification if necessary. Finally, we interpolated the missing electrodes and re-referenced the data to average reference.

For TMS-EEG data, we first cut the data into 3.2 second epochs (−1.1 to +2.1 seconds from the first TMS pulse) and performed baseline correction using data from the fixation period (−1 to −0.6 seconds from the first TMS pulse). The large TMS-induced artefact for each TMS pulse was removed from -2 ms to 10 ms around each pulse by setting the data at these timepoints to zero, after which we down-sampled the data from 25000 Hz to 1000 Hz. We then identified and removed bad trials and channels and ran fastICA using the same way as described above. In the first round of ICA, we only removed TMS-evoked muscle artefacts with large amplitudes. We then extended data removal from -2 ms to 15 ms around each pulse and interpolated the data with cubic interpolation ^43,79^. Afterwards, we filtered the data with a band-pass filter (0.01-100 Hz) and a stop-band filter (48-52 Hz) using the Butterworth method. In the second round of ICA, we removed all the remaining artefacts reflecting eye movement, blink-related activity, electrode artefacts, and residual muscle activity. Finally, we interpolated the missing electrodes, re-referenced the data to average reference, and re-epoched the data to stimulus onset for further analysis.

### Measuring the individual alpha frequency (IAF)

In each session, we measured the pre-stimulus IAF in the first delay for each participant using the data of three EEG-only blocks by performing a fast Fourier transformation (*fft*) in Matlab. The pre-stimulus IAF was defined as the frequency with the maximum local power in the alpha band (8-13 Hz) over a cluster of posterior electrodes around the right IPS (P2, P4, P6, P8, PO4, PO8, O2) during the first delay (between -600 ms and stimuli onset) across conditions.

### Time-frequency decomposition and event-related potentials (ERP)

We analysed the effects of alpha rTMS entrainment on oscillatory power and ITPC using time-frequency decomposition. We applied a Morlet continuous wavelet transform (defined with *f* = 5, *σ* = 0.4) to each trial using custom Matlab scripts and then averaged the results across trials. This procedure was performed separately for each participant, condition and channel. To visualise the time-frequency data, we averaged over a cluster of right posterior electrodes (P2, P4, P6, P8, PO4, PO8, O2, CP2, CP4, CP6, TP8, C2, C4, C6, T8) and calculated the differences between rTMS and aTMS. We selected this relatively large region of interest due to the variability of the individualized TMS target across the right posterior areas. For the topographic maps, we calculated alpha power (8-13 Hz) for each electrode and averaged across the time window either before (−400 to 0 ms) or after stimulus onset (0 to 400 ms). For the time-frequency results with evoked activity removed (Figure S3), we subtracted the evoked potentials (averaged signal across trials for each condition separately) from each trial and then performed Morlet wavelet transform as described above.

For our analysis of posterior-contralateral ERP components, we subtracted the waveform from the posterior hemisphere ipsilateral to the cued location from the posterior hemisphere contralateral to the cued location. For this we used the same electrodes as in the time-frequency analysis, plus the equivalent electrodes in the left hemisphere. We subtracted the mean response in the baseline of -1400 to -1350 ms for the results shown in Figure 2 (while we subtracted the mean baseline activities of 100 ms period prior to stimulus onset as a comparison in Figure S4).

### EEG multivariate decoding analysis

We performed multivariate decoding analysis using the CosMoMVPA toolbox in Matlab ^80^. We used two classifiers, based on data reflecting evoked potentials and alpha power, on three types of information (*where to attend*: attend left vs. attend right; *what to attend to*: attend colour vs. attend orientation; *visual feature information*: yellowish vs. blueish for the colour task, leftward vs. rightward for the orientation task). For visual feature information, we decoded visual features in four attention conditions: the attended feature of the attended object (aOaF), the unattended feature of the attended object (aOuF), the attended feature of the uattended object (uOaF), and the unattended feature of the unattended object (uOuF). For example, for a trial cuing attend left and attend orientation, the aOaF was the orientation of the left object, and the classifier was required to decode between trials on which the left object was oriented leftward vs. rightward.

The preprocessed EEG data was further down-sampled to 50 Hz to reduce computation time and to improve the stability of decoding ^8^. For ERP-based decoding, we band-pass filtered the data to 1-6 Hz to isolate low-frequency evoked potentials as described in ^8^. To increase signal to noise, we then averaged pairs of trials into ‘pseudo-trials’ using a random selection of the available trials ^81^. We then trained and tested a linear support vector machine classifier (LibSVM), on the filtered instantaneous voltage across all sensors at each timepoint (the interval between two timepoints is 20 ms), to distinguish the category of the trials using leave-one-pseudo-trial-out cross-validation. We repeated this random averaging procedure 100 times per participant, and used the average over these 100 repetitions as the decoding accuracy for statistical testing, to ensure that the results were not dependent on a specific set of trial averages. For comparison with previous literature using the full instantaneous EEG voltage at each timepoint, we repeated the same procedure on data that were down-sampled to 150 Hz and filtered to between 1-40 Hz. This showed highly similar results to the ERP-based decoding (Figure S2).

The procedure for alpha-based decoding was the same as ERP-decoding, except that instead of low-pass filtering, we band-pass filtered the data to the alpha band (8-13 Hz) and then used the Hilbert transform (*hilbert* function in Matlab) to extract the complex spectrums to calculate instantaneous alpha power by squaring the alpha amplitude obtained from the *abs* function in Matlab. We then ran the decoding as above, using the alpha power time series at each channel. For the alpha-based decoding results after removing evoked potentials (Figure S5), we first subtracted the evoked potentials (averaged signal across trials for each condition separately) from each trial and then performed the same procedures described above. For comparison, we conducted power-based decoding based on a broad range of frequencies (5 Hz bands between 13 and 48 Hz; Figure S1) with sampling rate at 50 Hz (for 13-18 Hz band), 100 Hz (for 18-23 Hz, 23-28 Hz, and 28-33 Hz bands), or 150 Hz (for 33-38 Hz, 38-43 Hz, and 43-48 Hz bands). We did not include the theta range (4-8 Hz) as that would overlap with the ERP-based decoding (1-6Hz).

## Statistical analysis

For decoding analysis, we used Bayesian t-tests with bayesFactor toolbox in Matlab (https://github.com/klabhub/bayesFactor) to compare the decoding accuracy to chance at every time point and track the decoding dynamics (BF_10_ > 3 as strong evidence for H1; BF_10_ < 1/3 as strong evidence for H0). For this we used a half-Cauchy prior with an interval of 0.5 and a default width of 0.707 ^54^. We also performed repeated-measures Bayesian ANOVAs to compare the decoding results for different conditions in each time point for each decoding. For analysis of alpha entrainment and ERPs, we similarly implemented repeated-measures Bayesian ANOVAs in each time point to examine the rTMS effects on alpha power, ITPC, and ERPs. To evaluate the correlation between TMS-induced changes in behavioural performance and decoding accuracy of the information about *where to attend*, we used Kendall’s Tau to examine the rank correlation between TMS-induced changes in ERP- and alpha-based decoding, at each timepoint, and behavioural changes in response error and RT. Then we calculated one-sided Bayes Factor for each Kendall’s Tau, using the method proposed by ^56^. This test is one-sided because we pre-specified the direction that larger neural signatures of attention should predict better performance.

## Acknowledgements

This project was supported by UKRI MRC intramural funding (SUAG/093/G116768) to A.W., UKRI MRC award MC_PC_20046, and a Gates Cambridge Scholarship awarded to R.L. (OPP1144). For the purpose of open access, the author has applied a Creative Commons Attribution (CC BY) licence to any Author Accepted Manuscript version arising from this submission.

## Author Contributions

R.L., J.D., and A.W. conceived the project; R.L., E.M., J.D., and A.W. designed the experiment; R.L., E.M., J.B.J., and A.W. implemented the experiment; R.L. conducted the experiment; R.L., E.M., C.L.S., and A.W. analysed data; R.L. wrote the first draft of the paper; R.L., E.M., C.L.S., J.B.J., J.D., and A.W. contributed to the final draft of the paper; A.W. provided overall supervision; and A.W. and R.L contributed funding.

## Declaration of interests

The authors declare no competing interests.

---

**Figure S1.**
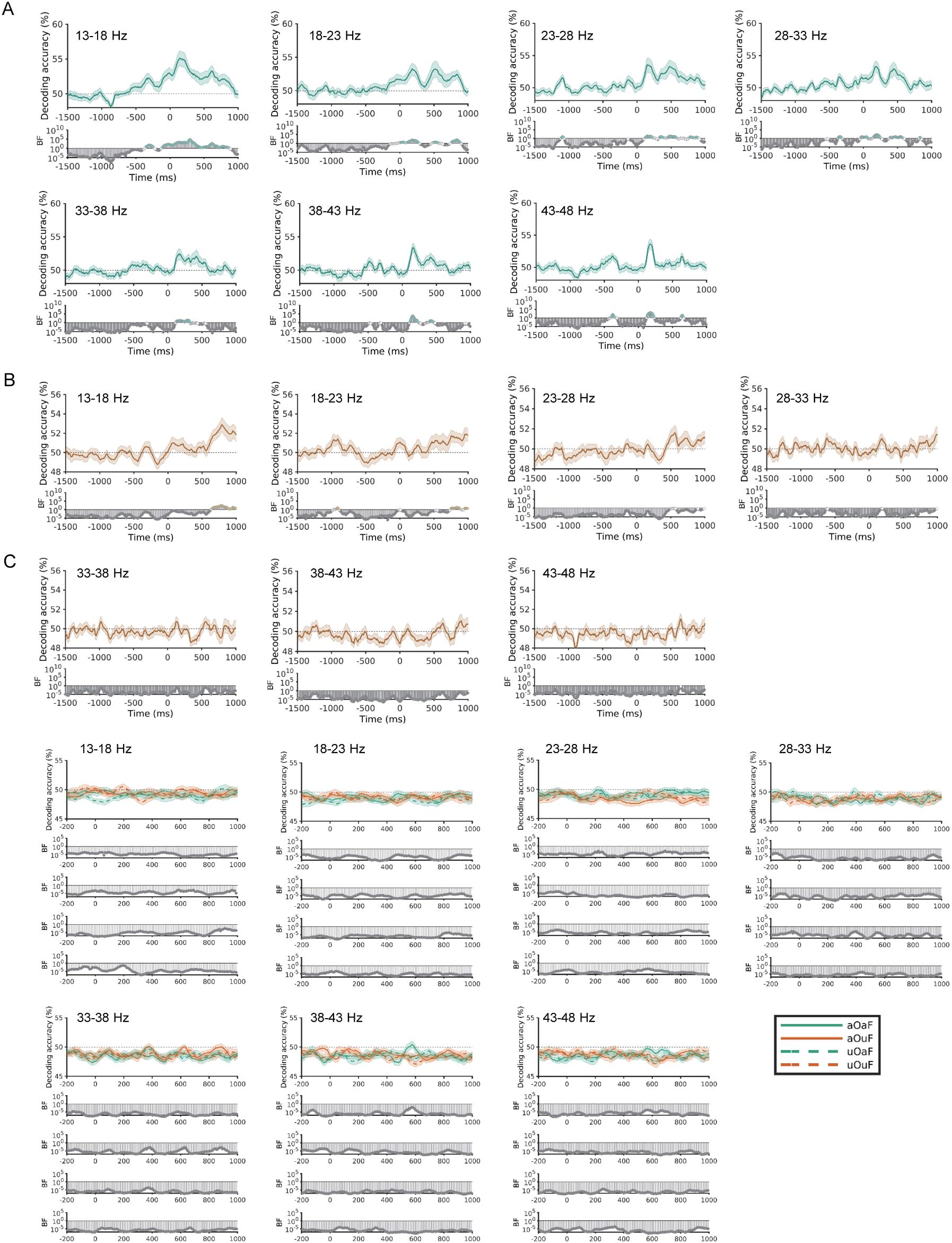
EEG decoding results of the information about *where to attend*, *what to attend to* and *visual features* based on power of different frequency bands using EEG-only data. EEG power-based decoding results of (A) *where to attend* (attend left vs. attend right), (B) *what to attend to* (attend colour vs. attend orientation), and (C) *visual feature information*. Time in all plots is relative to stimulus onset (ms). Shaded error bars indicate the standard error. The bottom rows of each plot show the Bayes Factors (BF_10_) at each time point on a log scale, with BF_10_ < 1/3 marked in grey and BF_10_ > 3 highlighted with coloured circles). aOaF: attended feature of attended object; aOuF: attended feature of unattended object; uOaF: unattended feature of attended object; uOuF: unattended feature of attended object.

**Figure S2.**
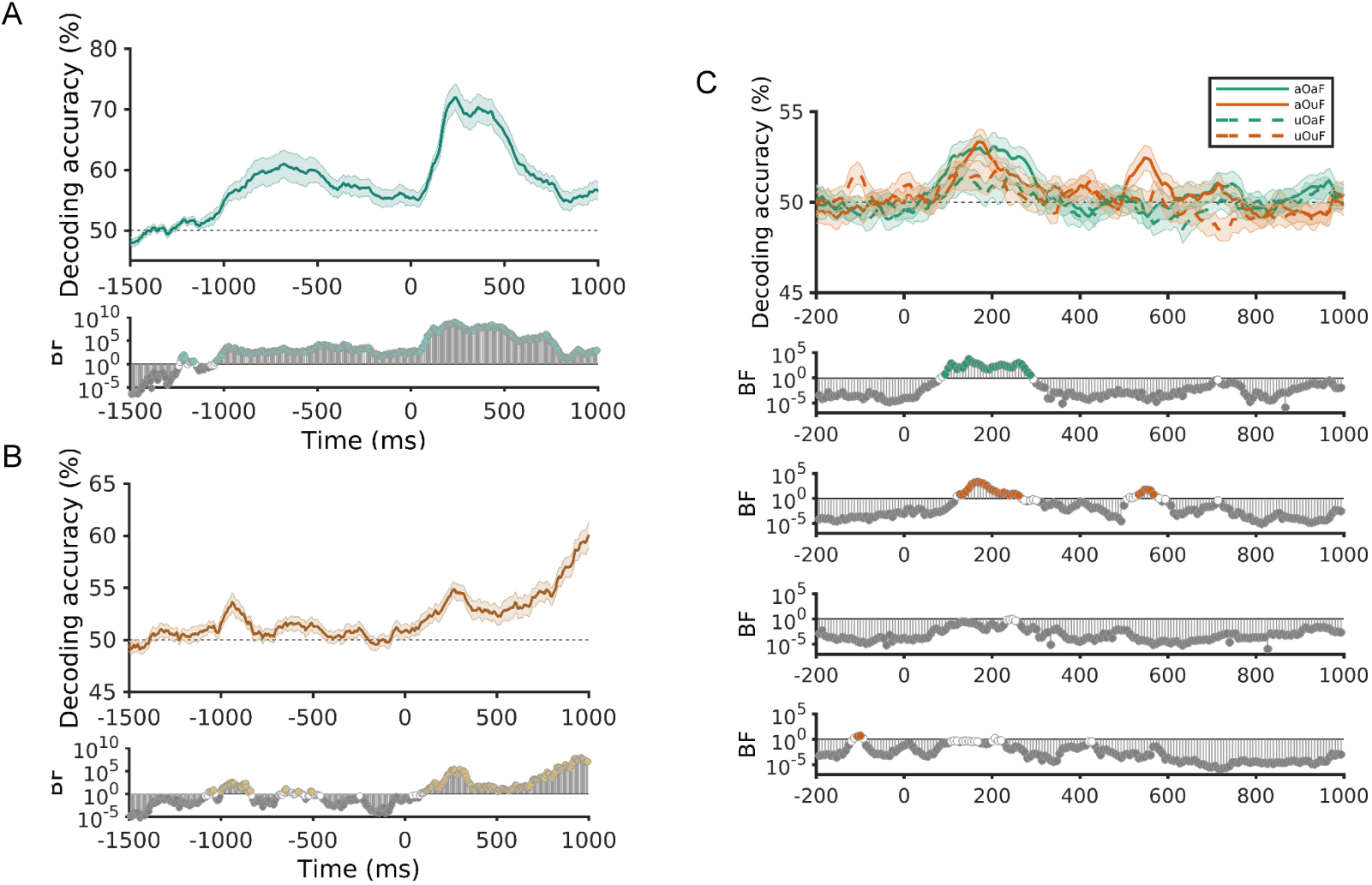
EEG voltage-based decoding results of the information about *where to attend*, *what to attend to* and *visual features* using EEG-only data. EEG voltage-based decoding (1-40 Hz) results of (A) *where to attend* (attend left vs. attend right), (B) *what to attend to* (attend colour vs. attend orientation), and (C) *visual feature information*. Time in all plots is relative to stimulus onset (ms). Shaded error bars indicate the standard error. The bottom rows of each plot show the Bayes Factors (BF) at each time point on a log scale, with BF_10_ < 1/3 marked in grey and BF_10_ > 3 highlighted with coloured circles). aOaF: attended feature of attended object; aOuF: attended feature of unattended object; uOaF: unattended feature of attended object; uOuF: unattended feature of attended object.

**Figure S3.**
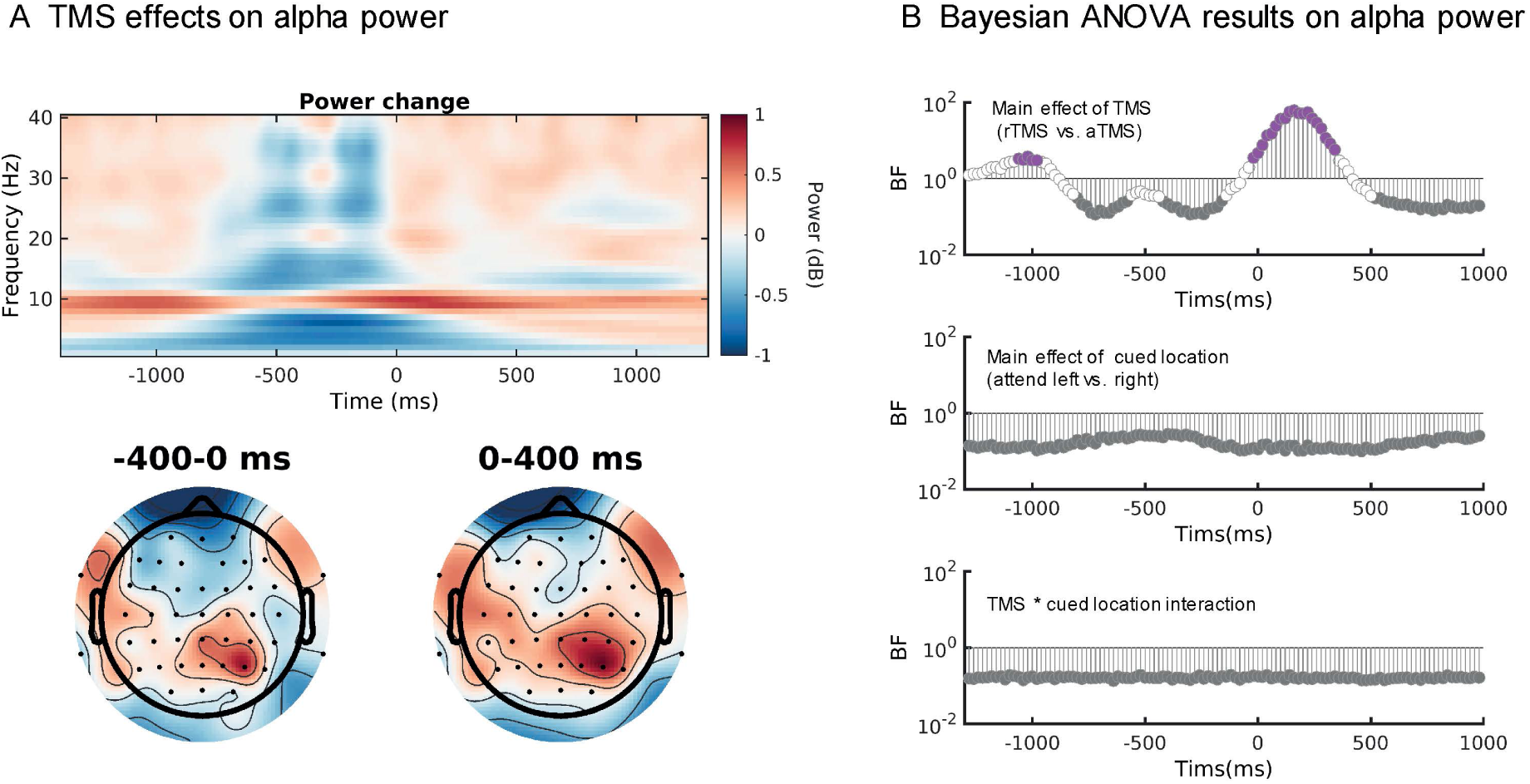
TMS entrained alpha oscillations after subtracting evoked activtiy. (A) Time-frequency analysis for alpha power entrainment (alpha rhythmic TMS – arrhythmic TMS; averaged across attend left and attend right conditions over right posterior electrodes). Evoked activity was subtracted from each trial for each condition to remove the potential phase-locked non-oscillatory components. The topographic plots show the scalp distribution of the alpha power entrainment effect in time windows before and after stimulus onset. (B) Results of repeated-measure Bayesian ANOVAs (TMS: rTMS vs. aTMS; cued location: attended left vs. right) for each timepoint on alpha power.

**Figure S4.**
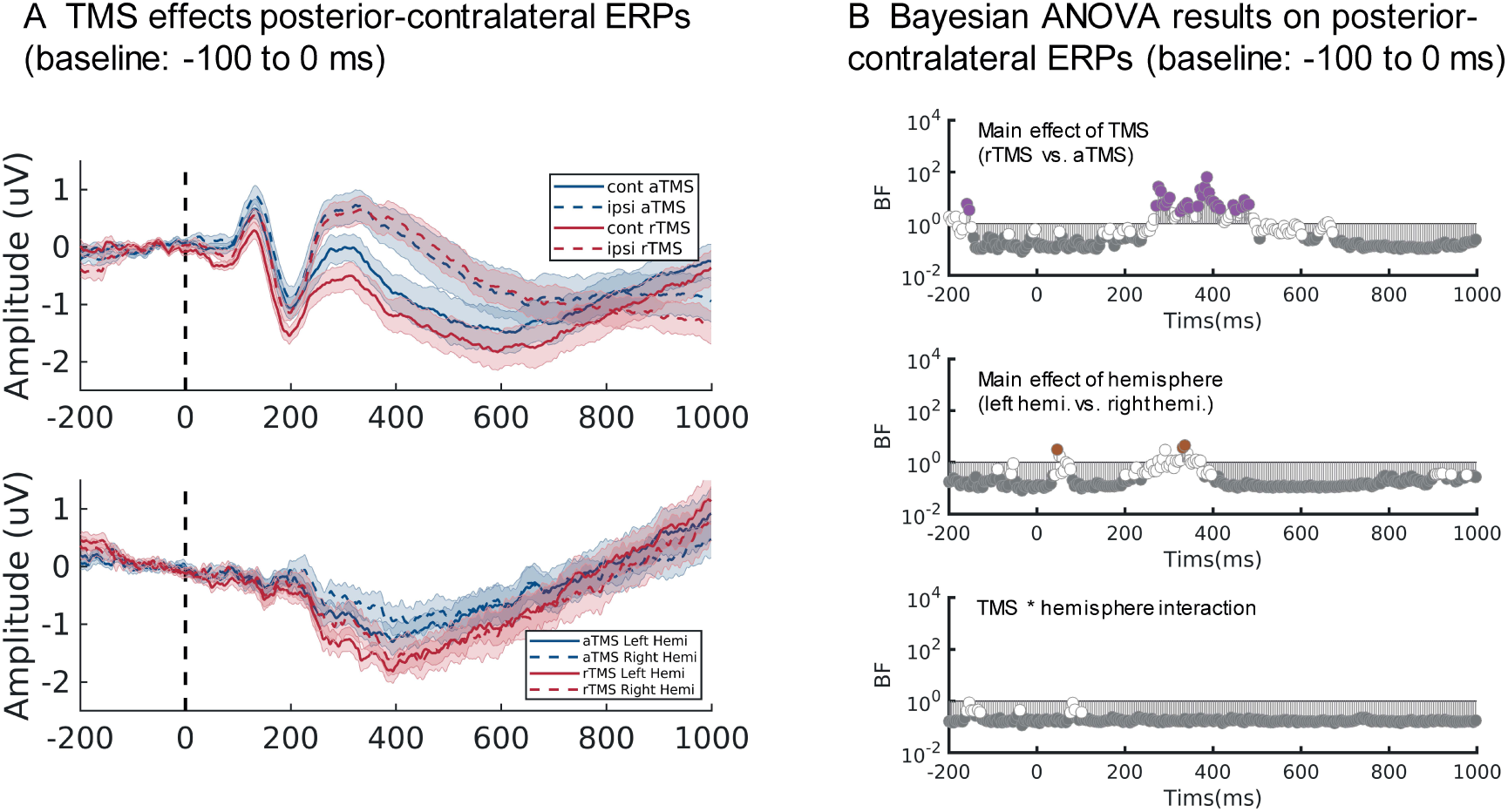
TMS effects on posterior-contralateral amplitudes with pre-stimulus baseline correction. (A) The upper plot shows the ERP waves in the contralateral vs. ipsilateral hemisphere to the spatial cue during different TMS protocols using posterior electrodes (see Methods). The lower plot shows the contralateral minus ipsilateral ERP waves in the left and right hemispheres separately. Baseline correction was performed using the averaged data between - 100 and 0 ms. Shaded error bars indicate the standard error. (B) Results of repeated-measure Bayesian ANOVAs (factors: TMS: rTMS vs. aTMS; hemisphere: left hemisphere vs. right hemisphere) for each timepoint on evoked potentials (contralateral – ipsilateral wave; the lower plot of Figure S4A). In all plots, time is relative to stimulus onset (ms). Bayes Factor (BF_10_) at each time point was on a log scale, and we marked the BF_10_ with BF_10_ < 1/3 in grey and BF_10_ > 3 in coloured circles. We downsampled the Bayesian ANOVAs results to 200 Hz in for visualisation.

**Figure S5.**
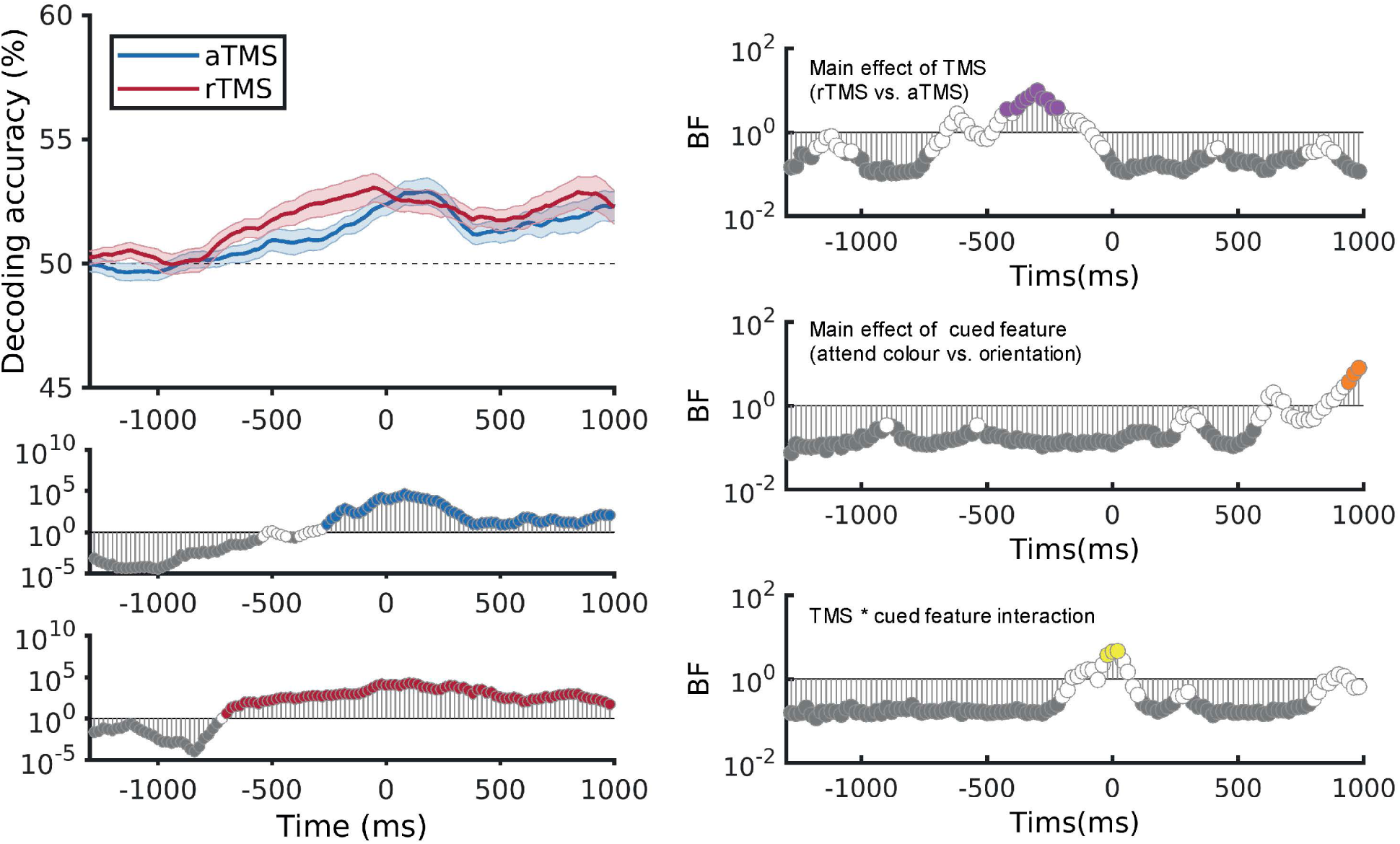
TMS effects on decoding accuracy of *where to attend* (attend left vs. attend right) based on alpha power with evoked activity removed. Left: TMS effects on decoding of *where to attend* (attend left vs. attend right) based on alpha power with evoked activity removed. The upper plot shows decoding accuracies, and the middle and lower plots show Bayes Factors (BF_10_) for a comparison of decoding accuracy against chance (50%) at each timepoint, for aTMS (middle, blue) and rTMS (bottom, red). Right: Results of repeated-measure Bayesian ANOVAs (TMS: rTMS vs. aTMS; attended feature: attend colour vs. attend orientation) for each timepoint on decoding accuracy based on alpha power. Time in all plots is relative to stimulus onset (ms). Cues were shown at approximately -1100 ms (adjusted for IAF). BF_10_ at each time point is shown on a log scale. We marked the BF_10_ with BF_10_ < 1/3 in grey and BF_10_ > 3 in coloured circles.

**Table S1.** Descriptive statistics: Mean and standard deviation (in brackets) of response error and reaction time (RT) in different TMS conditions.

|  |  | Cued left |  | Cued right |  |
| --- | --- | --- | --- | --- | --- |
|  |  | Cued colour | Cued orientation | Cued colour | Cued orientation |
| Error<br>(degrees) | No TMS (N = 31) | 10.44 (4.02) | 6.78 (1.92) | 9.13 (3.02) | 6.72 (1.84) |
|  | rTMS (N = 32) | 10.69 (6.06) | 7.27 (3.87) | 10.50 (5.70) | 6.78 (2.22) |
|  | aTMS (N = 32) | 10.82 (8.17) | 7.13 (3.04) | 10.76 (6.24) | 7.22 (3.01) |
| RT (s) | No TMS (N = 31) | 1.65 (0.41) | 2.09 (0.46) | 1.64 (0.40) | 2.10 (0.47) |
|  | rTMS (N = 32) | 1.49 (0.37) | 1.83 (0.52) | 1.50 (0.37) | 1.83 (0.52) |
|  | aTMS (N = 32) | 1.52 (0.37) | 1.88 (0.47) | 1.52 (0.37) | 1.89 (0.50) |

## Notes

### Competing Interest Statement

The authors have declared no competing interest.

### Summary of Updates

We added some more supplementary figures and changed the title.

